# Chemical structures of cyclic ADP ribose (cADPR) isomers and the molecular basis of their production and signaling

**DOI:** 10.1101/2022.05.07.491051

**Authors:** Mohammad K. Manik, Yun Shi, Sulin Li, Mark A. Zaydman, Neha Damaraju, Samuel Eastman, Thomas G. Smith, Weixi Gu, Veronika Masic, Tamim Mosaiab, James S. Weagley, Steven J. Hancock, Eduardo Vasquez, Lauren Hartley-Tassell, Natsumi Maruta, Bryan Y. J. Lim, Hayden Burdett, Michael J. Lansdberg, Mark A. Schembri, Ivan Prokes, Lijiang Song, Murray Grant, Aaron DiAntonio, Jeffrey D. Nanson, Ming Guo, Jeffrey Milbrandt, Thomas Ve, Bostjan Kobe

**Affiliations:** The University of Queensland, School of Chemistry and Molecular Biosciences and Australian Infectious Diseases Research Centre, Brisbane, QLD 4072, Australia; Institute for Glycomics, Griffith University, Southport, QLD 4222, Australia; Department of Pathology and Immunology, Washington University School of Medicine in St. Louis, MO 63100, USA; Department of Developmental Biology, Washington University School of Medicine in Saint Louis, St. Louis, MO 63100, USA; Department of Genetics, Washington University School of Medicine in Saint Louis, St. Louis, MO 63100, USA; Department of Plant Pathology, University of Nebraska-Lincoln, Lincoln, NE 68583, USA; Department of Chemistry, University of Nebraska-Lincoln, Lincoln, NE 68588, USA; Edison Family Center for Genome Sciences and Systems Biology, Washington University School of Medicine in St. Louis, MO 63110, USA; Department of Chemistry, University of Warwick, Coventry, CV4 7AL, UK; School of Life Sciences, University of Warwick, Coventry, CV4 7AL, UK; Department of Agriculture and Horticulture, University of Nebraska-Lincoln, Lincoln, NE 68583, USA; The University of Queensland, Institute for Molecular Bioscience, Brisbane, QLD 4072, Australia

## Abstract

Cyclic ADP ribose (cADPR) isomers are important signaling molecules produced by bacterial and plant Toll/interleukin-1 receptor (TIR) domains via NAD^+^ hydrolysis, yet their chemical structures are unknown. We show that v-cADPR (2’cADPR) and v2-cADPR (3’cADPR) isomers are cyclized by *O*-glycosidic bond formation between the ribose moieties in ADPR. Structures of v-cADPR (2’cADPR)-producing TIR domains reveal that conformational changes are required for the formation of the active assembly that resembles those of Toll-like receptor adaptor TIR domains, and mutagenesis data demonstrate that a conserved tryptophan is essential for cyclization. We show that v2-cADPR (3’cADPR) is a potent activator of ThsA effector proteins from Thoeris anti-phage defence systems and is responsible for suppression of plant immunity by the effector HopAM1. Collectively, our results define new enzymatic activities of TIR domains, reveal the molecular basis of cADPR isomer production, and establish v2-cADPR (3’cADPR) as an antiviral signaling molecule and an effector-mediated signaling molecule for plant immunity suppression.

**One-Sentence Summary:** The chemical structures of two *O*-glycosidic bond-containing cyclic ADP ribose isomers, the molecular basis of their production, and their function in antiviral and plant immunity suppression by bacteria are reported.

## Main Text

The ∼150-residue TIR (Toll/interleukin-1 receptor) domains are widely distributed in animals, plants and bacteria, and function through self-association and homotypic interactions with other TIR domains (*1*). In plants and animals, these domains are predominantly found in proteins with immune functions such as TLRs (Toll-like receptors), IL-1Rs (interleukin-1 receptors) and their adaptor proteins (*2–5*), and plant NLRs (nucleotide-binding, leucine-rich repeat receptors) (*6, 7*). TIR domains form higher-order oligomers and orchestrate signal amplification by a mechanism referred to as signaling by cooperative assembly formation (SCAF) (*6, 8–10*).

In bacteria, initial studies suggested that TIR domain-containing proteins, such as TlpA, TcpB, TcpC, TirS, PumA and TcpS, may serve as virulence factors by inhibiting host innate immune signaling by molecular mimicry (*11–20*). However, it remains unclear how bacterial TIR domain-containing proteins enter the cell (Cirl et al., 2008; Rana et al., 2013; Spear et al., 2009). More recently, bacterial TIR domain-containing proteins have been implicated in antiphage defence systems (*21–24*).

Many TIR domains have been found to cleave NAD^+^ (nicotinamide adenine dinucleotide) (*25–28*). In animals, SARM1 (sterile alpha and Toll/interleukin-1 receptor motif-containing 1) is a metabolic sensor of axonal NMN (nicotinamide mononucleotide) and NAD^+^ levels, and its TIR domain can execute programmed axon death by cleaving NAD^+^ into Nam (nicotinamide) and either ADPR (ADP-ribose) or cyclic ADPR (cADPR) (*25, 28, 29*). The TIR domains of plant NLRs (TNLs) similarly cleave NAD^+^ into Nam and either ADPR or a cADPR isomer known as variant cADPR (v-cADPR) (*26*), which has a different but unknown cyclization site, compared to canonical cADPR (*30*). A conserved catalytic glutamate is required for NAD^+^ hydrolysis by SARM1 and plant TIR domains, and the same glutamate is required for SARM1 to trigger axon degeneration and plant TNLs to trigger a localized cell death in response to infections (*25, 26, 28*). The v-cADPR isomer is a biomarker of plant TNL enzymatic activity, but may not be sufficient in itself to trigger a cell-death response in plants (*31*).

Bacterial TIR domains have also been found to be capable of cleaving NAD^+^ (*22, 27, 32–34*), again producing Nam and ADPR or the distinct cADPR isomers v-cADPR or v2-cADPR. In bacteria, NADase activity by TIR domains has recently been found to be a critical component of STING cyclic dinucleotide sensing (*22*) and the Thoeris defence system (*21*). The Thoeris system protects bacteria against phage infection and consists of two genes: *thsA* and *thsB* (the latter can be present in one or multiple copies). The first gene, *thsA,* encodes a protein that contains sirtuin-like or macro domain that bind NAD^+^ or its metabolites, and a SLOG-like domain suggested as a sensor for nucleotide-related ligands (*21, 35*). The second gene, *thsB*, encodes a TIR-domain-containing protein (ThsB) (*21, 34–37*). Upon phage infection, ThsB cleaves NAD^+^ and produces a cADPR isomer, which activates ThsA (*34*), causing further NAD^+^ depletion and cell death. A bacterial TIR-domain effector protein from *Pseudomonas syringae* DC3000, HopAM1, has further been shown recently to suppress plant immunity, by producing v2-cADPR (*33*). Although cADPR isomers have important immune and virulence functions, neither their chemical structures nor their specific mechanisms of action have been elucidated.

TIR-domain self-association is required for the NADase activity of SARM1 as well as plant TIR domains (*25, 28*) and recent crystal and cryo-EM (cryogenic electron microscopy) structures of SARM1 in its active conformation revealed that the active site spans two TIR-domain molecules, explaining the requirement of TIR domain self-association for NAD^+^ cleavage (*38*). The architectures of oligomeric TIR-domain assemblies formed by plant and SARM1 TIR domains are similar (*39, 40*) (termed “enzyme assemblies”), but are distinct from the assemblies formed by animal TIR domains involved in TLR signaling (termed “scaffold assemblies”) (reviewed by (*6*)). Both types of assemblies feature open-ended complexes with two strands of TIR domains in a head-to-tail arrangement, but differ in the orientation of the two strands (antiparallel in enzyme assemblies, parallel in scaffold assemblies). Some plant TIR domains have also been shown to act as 2′,3′-cAMP/cGMP synthetases, by hydrolyzing RNA/DNA, which requires a different TIR-domain oligomeric architecture than is required for NAD^+^ cleavage (*41*). In the case of bacterial TIR domains, a limited number of structures have been determined (*11, 12, 35, 42, 43*); the mechanism of NAD^+^ cleavage and cADPR isomer production, as well as the role of self-association in this process are poorly understood for any bacterial TIR domain.

In this study, we demonstrate that cADPR isomer-producing bacterial TIR domains can catalyze *O*-glycosidic bond formation between the ribose sugars in ADPR and that cyclization occurs between the anomeric position of the distal ribose and either the 2’ (v-cADPR; renamed here 2’cADPR) or the 3’ (v2-cADPR; renamed here 3’cADPR) position of the adenosine ribose. We report the cryo-electron microscopy (cryo-EM) structure of the filamentous assembly of AbTir^TIR^ in complex with a NAD^+^ mimic, which surprisingly resembles assemblies formed by TLR adaptors MAL and MyD88. Crystal structures of monomeric TIR domains highlight conformational changes associated with active assembly formation. We also show that a conserved tryptophan residue plays a key role in ADPR cyclization by bacterial TIR domains. We further demonstrate that v2-cADPR (3’cADPR) is a potent activator of ThsA from the Thoeris antiphage defence system, and present crystal structures of ThsA bound to v2-cADPR (3’cADPR) and in the ligand-free inactive state. These structures suggest that ThsA is activated by a change in its tetramer organization, which is induced by v2-cADPR (3’cADPR) binding to a conserved pocket in the SLOG domain. We finally show that v2-cADPR (3’cADPR) is responsible for suppression of plant immunity by the effector HopAM1.

## Results

### cADPR isomers have an *O*-glycosidic linkage between ribose sugars

To identify the sites of cyclization in cADPR isomers, we expressed and purified the TIR-domain regions of AbTir (AbTir^TIR^) and a TIR domain-containing protein from *Aquimarina amphilecti* (AaTir^TIR^; Fig. S1a), which have been reported to produce v- and v2-cADPR, respectively (*27, 44*). Comparative genomics analyses suggest that AbTir has unique functions distinct from the Thoeris defense system (Fig. S1b).

TLC (thin layer chromatography), HPLC (high-pressure liquid chromatography), real-time NMR-based and fluorescence-based NADase assays confirmed that purified AbTir^TIR^ and AaTir^TIR^ are enzymatically active and produce cADPR isomers (Fig. 1a, S1c-f). AbTir^TIR^-produced v-cADPR is identical to the cADPR isomer produced by TIR domains of the plant immune receptors L6 and ROQ1 (Fig. 1a). Both AbTir^TIR^ and AaTir^TIR^ can catalyze base-exchange reactions with 8-amino-isoquinoline (Fig. S2a). They can also use NADP^+^ as a substrate, but its cleavage only yields the products Nam and ADPPR, indicating that cADPR-isomer production is specific to NAD^+^ cleavage (Fig. S2b).

**Fig. 1.**
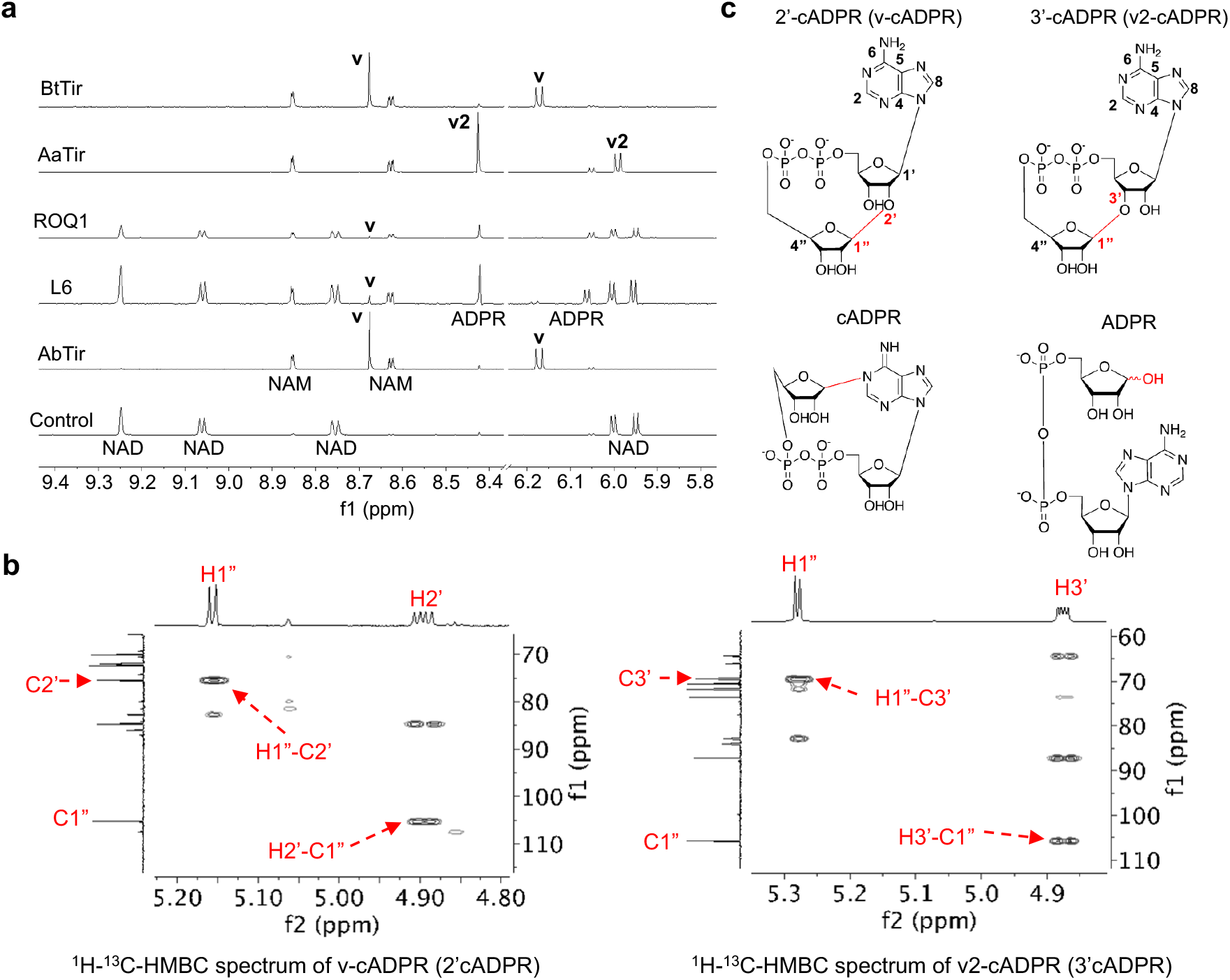
Chemical structures of v- and v2-cADPR. (a) Expansions of ^1^H NMR spectra showing NADase reactions by 0.1 μM AbTir^TIR^, 100 μM L6^TIR^, 26 μM ROQ1^TIR^, 0.5 μM AaTir^TIR^ and 2.5 μM BtTir^TIR^. The initial concentration for NAD^+^ was 500 μM, except for L6 it was 1 mM. Spectra correspond to 16 h incubation time, except for ROQ1 (incubation time 64 h). Selected peaks are labelled, showing the formation v-cADPR (**v**) and v2-cADPR (**v2**). (b) Expansions of HMBC spectra showing correlations through glycosidic linkages for 2’cADPR (v-cADPR) and 3’cADPR (v2-cADPR). (c) Chemical structures of 2’cADPR, 3’cADPR, cADPR and ADPR. Important NMR peaks (b) and their correlated positions in the chemical structure (c) are labelled, showing a 1”-2’ linkage for v-cADPR and a 1”-3’ linkage for v2-cADPR.

We purified v-cADPR and v2-cADPR from the reaction mixture, using HPLC, and determined their chemical structures using NMR (Fig. 1b, S2c-d, Table S1–2). The HMBC (heteronuclear multiple bond correlation) NMR spectrum of AbTir^TIR^-produced v-cADPR shows both H1’’-C2’ and H2’-C1’’ cross-peaks, revealing that AbTir^TIR^ can catalyze the formation of a ribose(1″→2′)ribose *O*-glycosidic linkage in ADPR (Fig. 1b). By contrast, HMBC spectrum of v2-cADPR, produced by AaTir^TIR^, shows both H1’’-C3’ and H3’-C1’’ cross-peaks, indicating that AaTir^TIR^ catalyzes the formation of a ribose(1″→3′)ribose *O*-glycosidic linkage in ADPR (Fig. 1b-c). Since the β-configuration of the anomeric carbon in NAD^+^ is retained in AbTir^TIR^ and AaTir^TIR^-catalyzed base-exchange reactions (Fig. S2a) and the coupling constants of the anomeric protons of v-cADPR (J1″,2″ ∼ 5.0 Hz) and v2-cADPR (J1″,3″ ∼ 4.4 Hz) are similar to those of NAD^+^ (5∼6 Hz), both cADPR isomers are likely to retain the same β-configuration as NAD^+^ and the base-exchange product. v2-cADPR purified from *N*. *benthamiana* leaves expressing the bacterial effector HopAM1 showed an identical chemical structure to that of AaTir^TIR^ (Table S3). Based on these chemical structures, we term the molecules 2’cADPR and 3’cADPR, highlighting the linkages between ribose rings of v-cADPR and v2-cADPR, respectively.

### Self-association enhances the NADase activity of cADPR isomer-producing TIR domains

SARM1 and plant TIR domains require self-association for their NADase activity (*25, 38–40*). We found that the NADase activity of AbTir^TIR^ increases disproportionally with increasing protein concentrations and increases substantially in the presence of molecular crowding agents (Fig. 2a-b). Furthermore, SEC-MALS (size-exclusion chromatography coupled with multiangle light scattering) experiments show that AbTir self-associates in a concentration-dependent manner (Fig. 3c-d), which requires both the TIR domain and the N-terminal coiled coil (CC) domain (AbTir^CC^). At high concentrations (100 µM), a construct encompassing both these domains (AbTir^full-length^) exists as a dimer in solution, whereas the TIR and CC domains both exist in a rapid monomer-dimer equilibrium (Fig. 2a-b). Interestingly, AbTir^full-length^ shows an initially suppressed enzymatic activity, followed by a sharp increase, suggesting that conformational rearrangements are required for efficient NAD^+^ cleavage by AbTir^full-length^ (Fig. S1c). Taken together, these results suggest that cADPR isomer-producing bacterial TIR domains may also require self-association for their NADase activity.

**Fig. 2.**
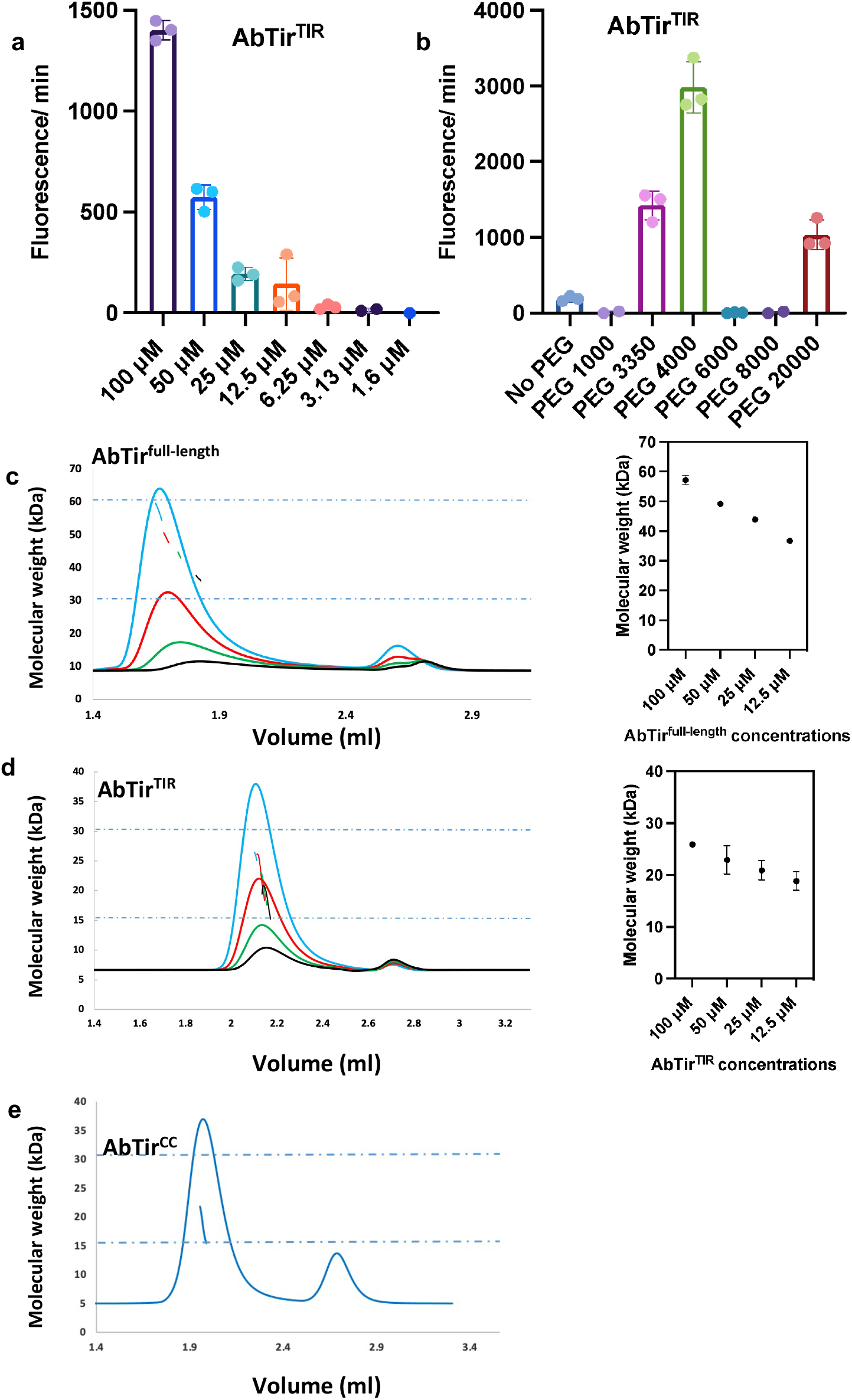
Self-association is vital for the enzymatic activity of AbTir. (a) NADase activity of AbTir^TIR^ at different concentrations, measured by the fluorescence assay using εNAD. (b) Effect of macromolecular crowding agents in the enzymatic activity, measured by the fluorescence assay using εNAD. In this experiment, 25 µM AbTir^TIR^ and 20% PEG were used. (c) Size-exclusion chromatography-coupled multi-angle light scattering (SEC-MALS) analysis of AbTir^full-length^ (1-269) (left panel). The elution of the protein from the SEC column (Superdex 200) was measured as a direct refractive index (dRI). SEC-MALS analysis of AbTir^TIR^ (134-269)on a Superdex 75 column (right panel). (d) Comparison of calculated and experimental molecular weight of the AbTir^full-length^ and its TIR domain.

**Fig. 3.**
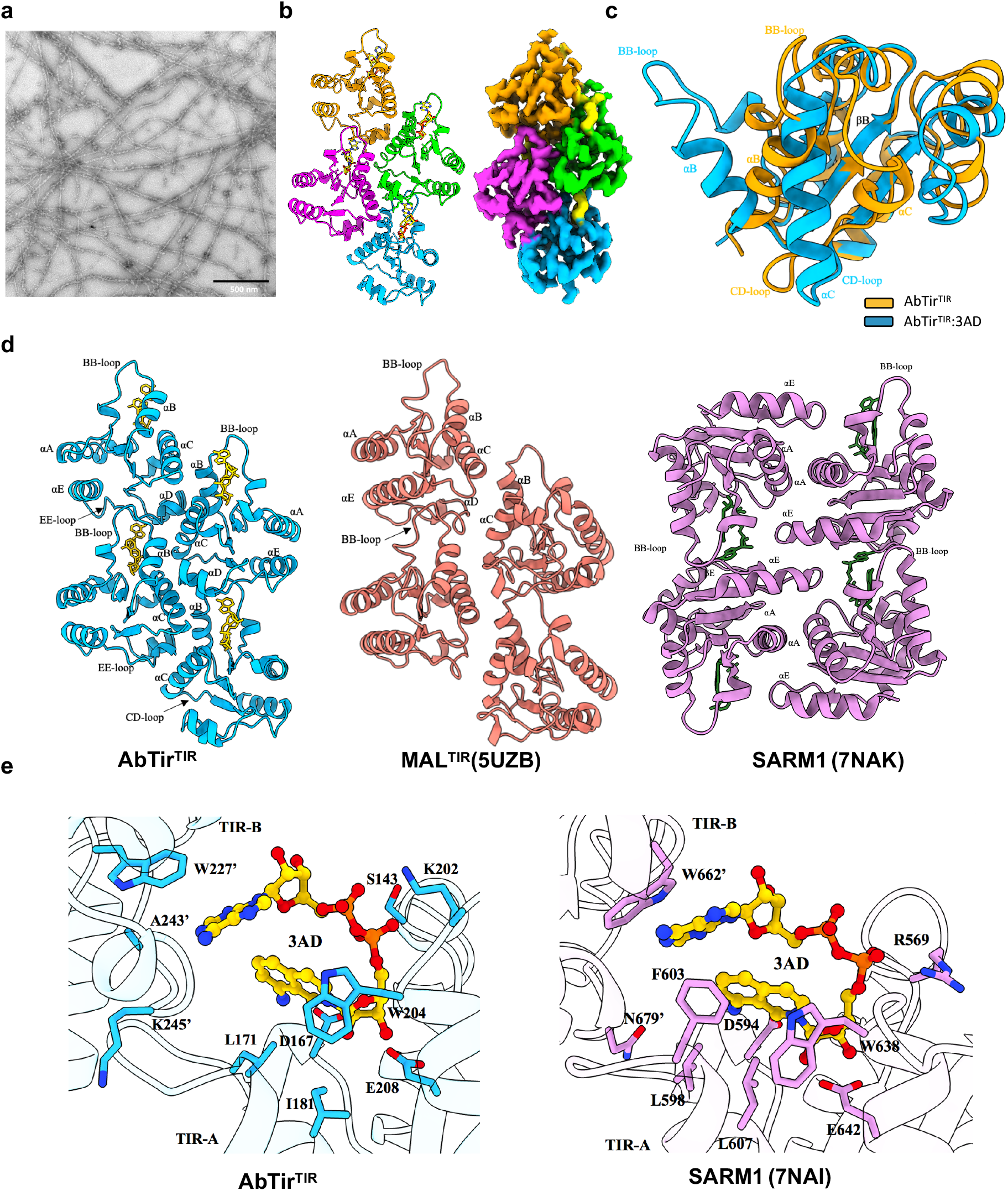
CryoEM structure of the AbTir^TIR^:3AD filament. (a) Negative-stain image of AbTir^TIR^:**3AD** filaments. (b) Cartoon representation and electrostatic potential density map of the AbTirTIR:**3AD** filament. TIR domain subunits are shown in blue, green, magenta and orange. **3AD** is shown in yellow. (c) Structural superposition of ligand-free (crystal structure, orange) and **3AD**-bound AbTir^TIR^ (cryo-EM structure, cyan) molecules reveal conformational changes in BB-loop and B and C helices upon substrate binding. (d) Comparison of AbTir^TIR^:**3AD**, MAL^TIR^ (PDB: 5UZB) and SARM1^TIR^:1AD (PDB: 7NAK) assemblies. (e) Comparison of the active sites in AbTir^TIR^ and the SARM1^TIR^ (PDB:7NAI) reveal similar substrate-binding modes.

### AbTir and BtTir TIR-domain crystal structures

To provide insights into the molecular basis of cADPR isomer production by TIR domains, we determined the crystal structures of AbTir^TIR^ and a closely related v-cADPR (2’cADPR)-producing TIR domain protein (47% sequence identity) from *Bacteroides thetaiotaomicron* (residues 156-287; BtTir^TIR^; Fig 1, S1a, and S2a), at 2.16 Å and 1.42 Å resolution, respectively (Table S4). Both structures have a canonical flavodoxin-like fold with a β-sheet comprised of five β-strands (βA-βE), surrounded by five α helixes (αA-αE) of variable lengths (the loops connecting the helices and strands are named according to the established nomenclature, e.g., the loop between βB and αB is known as the BB loop (*45*)) (Fig. S3a-b). The overall structure of AbTir^TIR^ closely resembles those of BtTir^TIR^, PdTir^TIR^ (PDB ID: 3H16) and the v-cADPR (2’cADPR)-producing TcpB^TIR^ (PDBs: 4C7M, 4LQC, 4LZP) (*11, 12, 42, 43*), with Cα RMSD (root-mean-square-distance) values of 1.2 Å, 1.8 Å and 1.9 Å, respectively (Fig. S3b-c). Both AbTir^TIR^ and BtTir^TIR^ show less similarity to the cADPR isomer-producing ThsB TIR domain of the *Bacilus cereus* MSX-D12 Thoeris defence system (BcThsB; PDB 6LHY; Cα RMSD values of 4.4 and 2.5 Å for AbTir^TIR^ and BtTir^TIR^, respectively). Significant differences are observed when comparing the CD-loop regions of AbTir, BtTir, PdTir and TcpB (Fig. S3b).

In the TIR domains with NADase activity (from SARM1, plant TNLs RPP1, ROQ1, and RUN1, and the oyster TIR-STING), the conserved glutamate residue essential for NADase activity is localized in a pocket consisting of residues from the βA strand, the AA and BB loops, and the αB and αC helices (*22, 25, 38, 39*). In AbTir^TIR^ and BtTir^TIR^, the αB and αC helices adopt significantly different conformations and the equivalent glutamate residue (E208 in AbTir and E230 in BtTir) is not located in a pocket, but is surface-exposed (Fig. S3d). Comparing the four individual chains in the asymmetric unit of the AbTir^TIR^ crystal structure, the region around E208 is highly flexible. Crystal packing of AbTir^TIR^ and BtTir^TIR^ reveals a common symmetric interface (Fig. S3c, S4a), with a large buried surface area (BSA; AbTir^TIR^, 1875 Å^2^; BtTir^TIR^, 1332 Å^2^), also observed in the crystal structures of PdTIR and TcpB (Fig. S4a). The structures suggest they represent an inactive conformation, possibly stabilized by the symmetric dimeric arrangement.

### Cryo-EM structure of AbTir^TIR^ bound to NAD^+^ mimic

We reasoned that we could capture the active state of AbTir by using base-exchange products of NAD^+^ hydrolysis, which are more biochemically stable than NAD^+^ itself and could resist cleavage at high concentrations of the enzyme (*38*). Indeed, in the presence of **3AD** (8-amino-isoquinoline adenine dinucleotide) we could visualize filamentous structures of AbTir^TIR^ by negative-stain electron microscopy (Fig. 3a). We collected cryo-EM data and determined the structure of these filaments at 3.4 Å resolution (Fig. S5, Table S5). The reconstruction clearly shows the presence of the **3AD** molecule between two monomers of AbTir^TIR^. Surprisingly, the structure reveals an arrangement of TIR domains different from the enzyme assemblies of SARM1 and plant TIR domains, but analogous to the scaffold assemblies formed by MAL and MyD88, each of which contains two parallel strands of TIR domains arranged head-to-tail (Fig. 3b, d, and S4b) (*6*). The intrastrand BE interface (BSA 710 Å^2^) involves the BB-loop, while the interstrand BCD interfaces involves one molecule interacting with two molecules in the parallel strand (BSA 1340 and 1410 Å^2^, respectively). Assembly formation is accompanied by remarkable conformational changes involving the BB-loop and the αB and αC helices (RMSD 1.8 Å for 101 Cα atoms; Fig. 3c, Movie S1). The BB-loop and αB helix have tilted outwards by ∼50-55° while the αC helix has refolded and includes residues from both the CC and CD loops. Despite the differences, the AbTir^TIR^ active site is very similar to that of SARM1^TIR^, including the conformation of the **3AD** molecule (Fig. 3e); the active site is formed by two TIR domains arranged through the analogous BE interface. The aromatic ring of the 8-amino isoquinoline group engages in interactions with L177 (αB helix) and W204 (αC helix) of AbTIR^TIR-A^, and the isoquinoline ribose is located in the cleft between the BB loop and the αB and αC helices with the C-2 and C-3 hydroxyls close to the key catalytic residue E208. The diphosphate group is involved in hydrogen-bonding interactions with the backbone amide of S143 and the sidechain of K202 in AbTIR^TIR-A^, while the adenine group forms interactions with the sidechains of W227 and the backbone A243 and K245 in AbTIR^TIR-B^. Similar to SARM1, the adenine-linked ribose is not involved in any direct interactions with either of the chains of the active site.

### Structure-guided mutagenesis reveals residues important for AbTir NADase activity

Using site directed mutagenesis, we verified the importance of residues in both the active site region and in the inter- and intrastrand interfaces, for NADase activity of AbTir^TIR^. The active site mutant E208A is completely inactive, while W204A, T205A and E208D have reduced NAD^+^ activity (Fig. 4a, b). Among the BB-loop mutants, D175A and L177A remain active, while G174A and S176A, show no or very low activity. Both of these residues are directly involved in intrastrand interactions suggesting that an intact BB loop is required for NADase activity. The interstrand interface mutant D182A has reduced activity, while R178A and Y207A are inactive, suggesting that active site stabilization by interstrand interactions is important for NADase activity. (Fig. 4a, b).

**Fig. 4.**
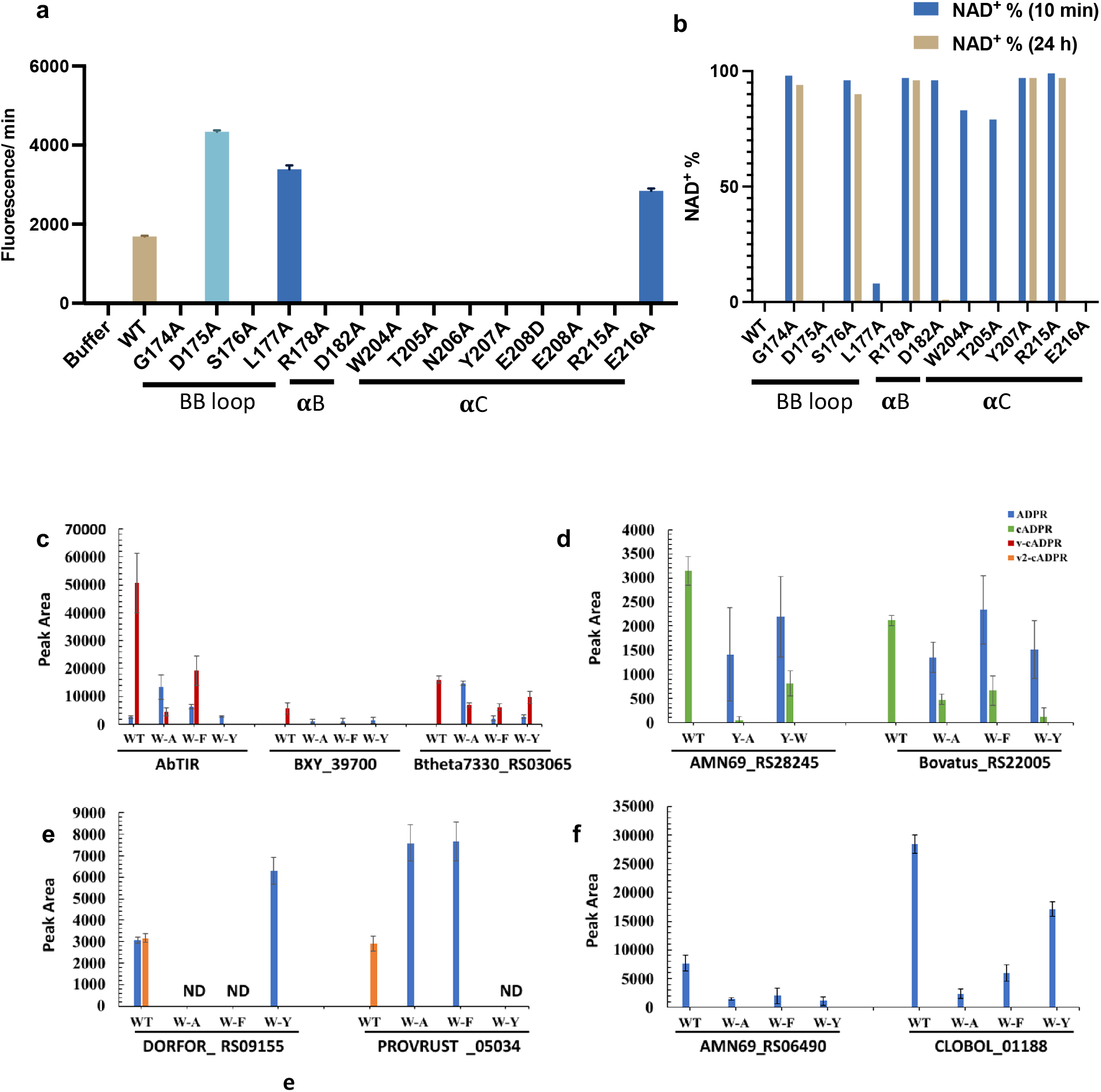
Mutagenesis of AbTir^TIR^ and conserved tryptophan in αC helix. (a) NADase activity of the different AbTir^TIR^ mutants, using the fluorescence-based assay, with 100 µM εNAD and 100 µM protein. Data are presented as mean ± SD (n = 3). (b) NAD^+^-cleavage activity of AbTir^TIR^ mutants, monitored by ^1^H NMR, using 500 µM NAD^+^ and 50 µM protein (except for the R178A mutant, where 30 µM protein and 300 µM NAD^+^ was used). (c-f) Mutations of the position equivalent to AbTir W204 affect the production of cyclic NAD^+^ catabolites by TIR domains. The NAD^+^ catabolite peak areas for wild-type and mutant TIR domain reactions after 1 hour are shown. TIR domain grouping is based on primary product of the wild-type protein (n=3 for all groups except where no data (ND) could be collected).

### Conserved αC helix tryptophan is required for ADPR cyclization

The products of NADase reaction were also assessed for the AbTir mutants (Table S6). The analysis revealed that only the W204A mutant has significantly reduced production of 2’cADPR (24%), compared to wild-type AbTir (93%), demonstrating its importance for ADPR cyclization (Table S6 and Fig. S6a). Next, we tested whether the mutation of the equivalent tryptophan residue has a similar effect on ADPR cyclization by other bacterial TIR domain-containing proteins. We analyzed the conservation of W204 in a multiple sequence alignment consisting of 122 previously functionally characterized TIR domains (*27, 28, 44*). We observed conservation of aromatic residues at this position among all TIR domains that produced a cyclic ADPR product (cADPR, v-cADPR [2’cADPR], or v2-cADPR [3’cADPR]) *in vitro*, with a strong preference for tryptophan - W (9/12), Y (2/12) and F (1/12) (Fig. S6b). This conservation is weaker among TIR domains that produce either non-cyclic products (Nam and ADPR) or lack activity altogether *in vitro*. These observations suggest that a large and aromatic residue at this position may play an important role in producing a cyclized ADPR product.

To further identify positions important for determining the product specificity of TIR domain NADases, we analyzed 278 TIR-domain sequences, and confirmed that the position corresponding to the tryptophan differs between cyclase and non-cyclase TIR domains, although not among cyclase TIR domains producing different forms of cADPR. All cyclase TIRs contain an aromatic residue at this position (Fig. S7a-e).

We tested the functional importance of this conserved aromatic residue by performing site-directed mutagenesis on TIR domains known to produce different products - v-cADPR (2’cADPR; AbTir, BXY39700, Btheta7330_RS03065), cADPR (Bovatus_RS22005, AMN69_RS28245), v2-cADPR (3’cADPR; ORFOR_RS09155, PROVRUST_05034), and ADPR (AMN69_RS06490, CLOBOL_01188). Mutations were designed to be conservative (W, Y or F) or disruptive (A). Reaction products were analyzed by HPLC (Fig. S6c-d). For all TIR domains producing a cyclic product (v-cADPR [2’cADPR], cADPR or v2-cADPR [3’cADPR]), the mutated proteins exhibited a decrease in the peak area corresponding to the cyclic product and an increase in the peak area corresponding to ADPR, when compared to the wild-type proteins (Fig. 4c-f). The non-conservative alanine mutations typically exhibited the greatest impact on the relative production of a cyclic product versus ADPR; however, this trend was not universally true. Mutations in the background of an ADPR-producing template had variable impacts on NADase activity (Fig. S6e), but ADPR always remained the only catabolite produced. These results are consistent with the hypothesis that the position equivalent to W204 in AbTir is critically important for the production of cyclic ADPR product.

### v2-cADPR (3’cADPR) is a potent activator of the Thoeris ThsA protein

Bacterial lysates containing cADPR isomers produced by bacterial ThsB or plant TIR domains have recently been shown to activate the ThsA NADase (Fig. 5a) of the *Bacillus cereus* MSX-D12 Thoeris antiphage defence system (*34*). To test if our purified cADPR isomers can directly activate ThsA, we produced and purified 4 different ThsA proteins (37-46% sequence identity) from *Bacilus cereus* MSX-D12 (BcThsA), *Acinetobacter baumannii* (AbThsA), *Enterococcus faecium* (EfThsA) and *Streptococcus equi* (SeThsA), and monitored the NAD^+^-cleavage activity of each in the absence and presence of ADPR, cADPR, v-cADPR (2’cADPR) and v2-cADPR (3’cADPR), using our NMR-based NADase assay (Fig. 5b-c and Fig. S8a-b). AbThsA, BcThsA and a SIR2 domain-only construct of BcThsA (BcThsA^SIR2^) rapidly cleave NAD^+^ (Fig. S8a-c), while EfThsA, SeThsA and SeThsA^SIR2^ (Fig. 5b-c and S8c) are almost inactive under the same conditions, with less than 10% of NAD^+^ consumed after 40 h. Neither BcThsA and AbThsA are further activated by ADPR, cADPR, v-cADPR (2’cADPR) or v2-cADPR (3’cADPR) (Fig. S8b), suggesting that these two proteins have been produced in a fully activated state. EfThsA and SeThsA rapidly cleave NAD^+^ in the presence of 500 μM v-cADPR (2’cADPR) or v2-cADPR (3’cADPR) (Fig. 5b-c). A clear dose-response is observed with v2-cADPR (3’cADPR) treatment and both proteins are strongly activated by v2-cADPR (3’cADPR) concentrations as low as 5 μM (Fig. 5c). In comparison, v-cADPR (2’cADPR) only activates these proteins at the highest concentration tested (500 μM). As expected, ADPR and cADPR have no effect on the NADase activity of EfThsA and SeThsA (Fig. 5b).

**Fig. 5.**
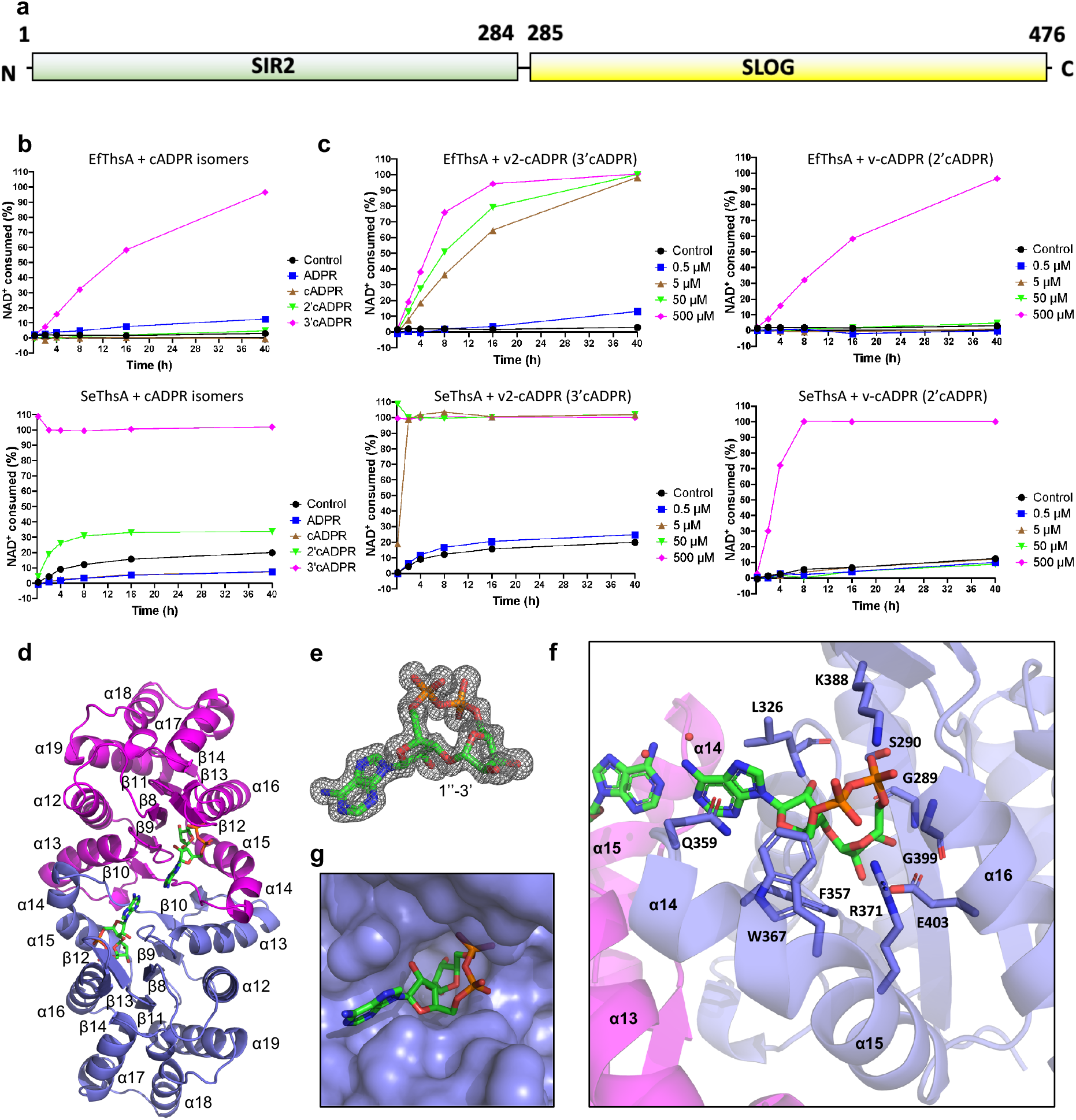
Thoeris ThsA is activated by v2-cADPR (3’cADPR) binding to a conserved pocket in its SLOG domain. (a) Schematic diagram of ThsA domain organization. Residue numbering corresponds to BcThsA. (b) Activation of EfThsA (0.5 μM) and SeThsA (10 μM) NADase activity by 500 µM ADPR, cADPR, v-cADPR (2’cADPR) and v2-cADPR (3’cADPR). The initial NAD ^+^ concentration was 500 µM. (c) Activation of EfThsA (0.5 μM) and SeThsA (10 μM) NADase activity by 0.5, 5, 50 and 500 µM v-cADPR (2’cADPR) and v2-cADPR (3’cADPR). The initial NAD ^+^ concentration was 500 µM. (d) Crystal structure of BcThsA^SLOG^ dimer (cartoon; chains coloured in slate and magenta) in complex with v2-cADPR (3’cADPR) (green stick). (e) Standard omit mFo-DFc map of v2-cADPR (3’cADPR), contoured at 3.0 σ. (f) Enlarged cutaway of the v2-cADPR (3’cADPR) binding pocket in the BcThsA^SLOG^ structure. (g) Surface representation of v2-cADPR (3’cADPR) binding pocket.

ITC (isothermal titration calorimetry) measurements showed that v2-cADPR (3’cADPR) binds directly to inactive (EfThsA) and activated (AbThsA) forms of ThsA at a ∼1:1 molar ratio, with *K_d_* values of 59.1 ± 15.8 nM and 189 ± 1.6 nM, respectively (Fig. S8d-e). No binding was detected for v-cADPR (2’cADPR) using ITC (Fig. S8d-e) and the weaker binding affinity of v-cADPR (2’cADPR) was also corroborated by competition binding assays via STD (saturation-transfer difference) NMR, as v2-cADPR (3’cADPR) almost eliminated or significantly reduced v-cADPR (2’cADPR) binding to EfThsA, SeThsA, BcThsA, and AbThsA at an equal concentration (Fig. S8f). Taken together, these findings support the model that ThsA is activated by TIR domain-produced cADPR isomers and demonstrate a strong preference for v2-cADPR (3’cADPR) over v-cADPR (2’cADPR).

### v2-cADPR (3’cADPR) binds to a highly conserved pocket in the ThsA SLOG domain

To provide structural insights into cADPR isomer selectivity by ThsA, we determined a crystal structure of the SLOG domain of BcThsA (BcThsA^SLOG^) in complex with v2-cADPR (3’cADPR) to 1.6 Å resolution (Fig. 5d, Table S4). Continuous electron density for a cADPR isomer with a ribose(1″→3′)ribose *O*-glycosidic linkage was observed, confirming the structural configuration assigned by our NMR assays (Fig. 5e). BcThsA^SLOG^ exists as a stable dimer in solution (Fig. S9, Table S7) and forms a symmetric dimer (SLOG dimer) in the crystal, with an identical interface to the dimer observed in the previously reported ligand-free BcThsA structure (Fig. 5d and S10a; PDB: 6LHX) (*35*). Binding of v2-cADPR (3’cADPR) does not lead to substantial structural rearrangements in either the SLOG domain or the dimer interface (Fig. S10a). v2-cADPR (3’cADPR) binds to a highly conserved pocket adjacent to the symmetric dimer interface and the adenine bases of the two v2-cADPR (3’cADPR) molecules in the dimer are only separated by 4.5 Å and bridged by two water molecules (Fig. 5f, S10b and Table S8). The C-2 and C-3 hydroxyls of the distal ribose of v2-cADPR (3’cADPR) interact with E403, while the diphosphate group is involved in hydrogen bonding interactions with S290, R371, K388 and the backbone amide and carbonyl of G289 and G399, respectively. The adenine base stacks against the side chains of L326 and Q359, while the C-2 hydroxyl of the adenine linked ribose forms a hydrogen bond with the backbone amide of L326. v-cADPR (2’cADPR), which has a ribose(1″→2′)ribose *O*-glycosidic linkage, cannot form this latter hydrogen bond and the adenosine moiety of this cADPR isomer is also likely to encounter steric hindrance with binding pocket residues (Fig. 5g), explaining the preference for 3’cADPR. Mutational analysis confirmed the importance of binding pocket residues for ThsA activation (Fig. S8g).

### v2-cADPR (3’cADPR) induces a change in ThsA tetramer organization

AbThsA, BcThsA, EfThsA, and SeThsA exist as tetramers in solution, and activation of SeThsA by v2-cADPR (3’cADPR) does not lead to a change in its oligomerization state (Fig. S9 and Table S7). However, the inactive SeThsA^SIR2^ exists as a dimer in solution, while the fully active BcThsA^SIR2^ exists as a monomer, suggesting that destabilization of SIR2:SIR2 domain interactions within the tetramer may be required for triggering the ThsA NADase activity (Fig. S8c, S9 and Table S7). To provide more detailed insight into how ThsA is activated by v2-cADPR (3’cADPR), we determined a ligand-free crystal structure of inactive SeThsA to 3.4 Å resolution (Table S4), and compared it to the published crystal structure of autoactive BcThsA (PDB: 6LHX) (*35*) and our BcThsA^SLOG^:v2-cADPR (3’cADPR) complex. Crystal packing analyses reveal a D2 symmetric tetramer with a core consisting of two SIR2 dimers flanked by SLOG dimers at both sides (Fig. 6a). Both SIR2 dimer interfaces involve residues from the α3, α7, α9 and α10 helices. The crystal structure of autoactive BcThsA (PDB: 6LHX) (*35*) has a tetramer with an identical architecture to SeThsA, but there are significant differences in the SIR2 dimer interfaces (Fig. 6a-b). One of the molecules in the BcThsA SIR2 dimers has undergone a rotation and translation of ∼25° and ∼14 Å compared to the SeThsA SIR2 dimers (Fig. 6b-c), resulting in a significant decrease of the interface area (2901.8 Å^2^ in SeThsA; 1334.8 Å^2^ in BcThsA). The α3 helix, which is involved in SIR2 dimerization, but also covers a part of the predicted active site region in SeThsA (Fig. 6b,d), adopts a different conformation (SIR2^A^ and SIR2^C^) or is disordered (SIR2^B^ and SIR2^C^) in BcThsA, enabling better access to catalytically important asparagine and histidine residues (N113/H153 in SeThsA; N112/H152 in BcThsA) (Fig. 6b,d) (*35*). Comparison of the SLOG dimers in the inactive SeThsA structure with the dimer in the BcThsA^SLOG^:v2-cADPR (3’cADPR) complex reveals that v2-cADPR binding is likely to induce significant changes in the orientation and position of the two SLOG domains (Fig. 6e-f; Movie S2). In the SeThsA tetramer, this change causes the SIR2^A^ and SIR2^B^, and the SIR2^C^ and SIR2^D^ domains to move in opposite directions, bringing the α10 helices into closer proximity (Fig. 6g, Movie S3), adopting a similar configuration to the SIR2 dimer interface observed in the crystal structure of autoactive BcThsA (Fig. 6b). We predict that this movement of the SIR2 domains is sufficient to destabilize the α3 helix conformation, enabling NAD^+^ to access the active sites. SeThsA double mutants with reverse-charge substitutions of highly conserved residues in the SIR2 dimer interface (Fig S10c) are either autoactive (E170R, D251R) or not activated by v2-cADPR (3’cADPR) (R166E, R254E), confirming the essential role of the SIR2 dimer interface in regulating ThsA NADase activity (Fig. 6h). In summary, these findings reveal the structural basis of cADPR isomer selectivity by ThsA and demonstrate that v2-cADPR (3’cADPR) activates the NADase function of ThsA by changing its tetramer organization.

**Fig. 6.**
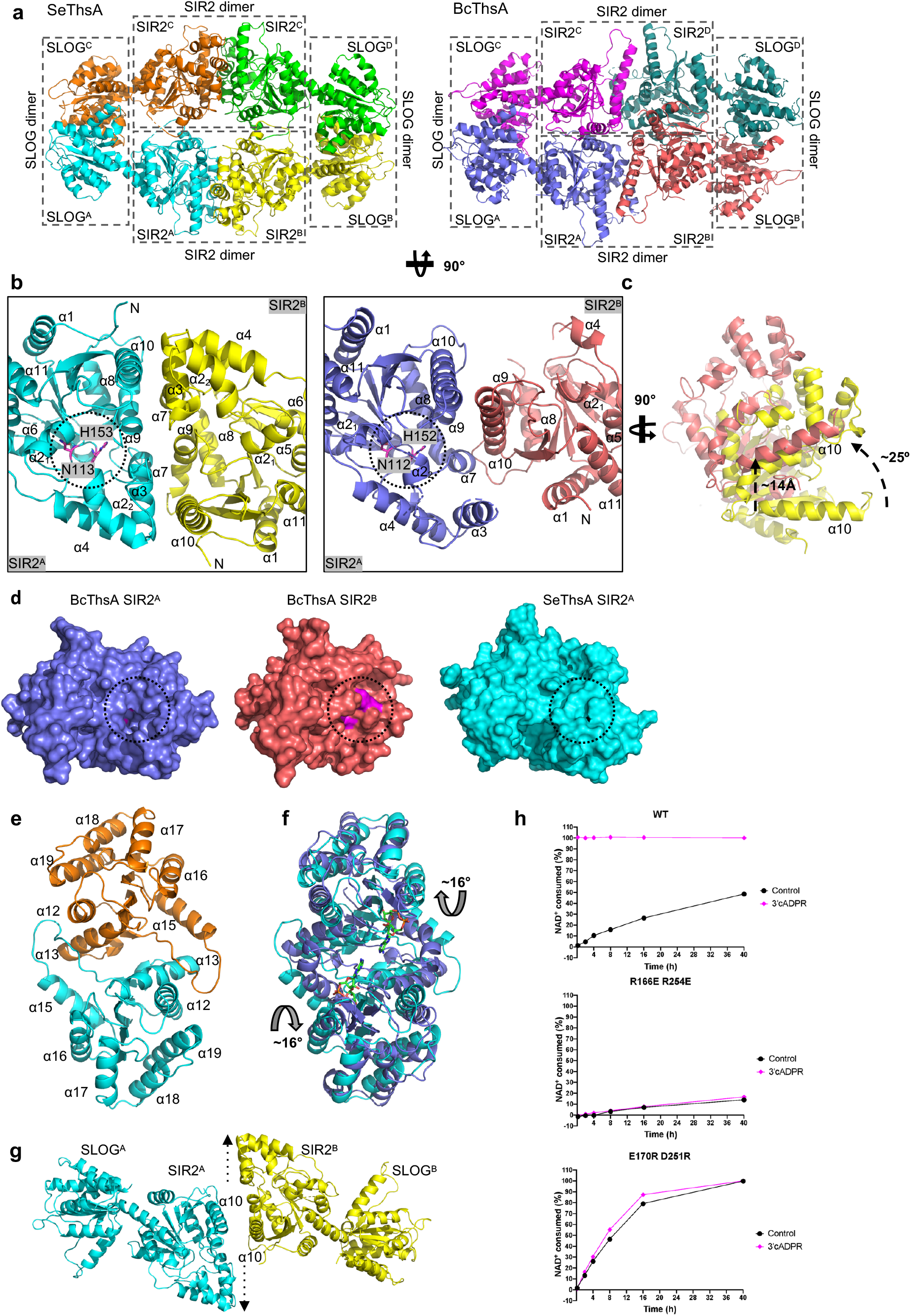
v2-cADPR (3’cADPR) induces changes to ThsA tetramer organization. (a) Structure of SeThsA (left; chains coloured in cyan, yellow, orange and green) and BcThsA (right; chains coloured in slate, salmon, magenta and teal) tetramers. SLOG and SIR dimers are highlighted by broken black boxes. (b) Enlarged cutaways of SeThsA (left) and BcThsA SIR2 dimers (right). Broken circles represent active site region with catalytically important residues (N113/H153 in SeThsA; N112/H152 in BcThsA) displayed as sticks (magenta). SIR2 dimers were superimposed using SIR2^A^ of SeThsA and BcThsA. (c) Comparison of SIR2^B^ in the superimposed SeThsA and BcThsA SIR2 dimers. Movement of BcThsA SIR2^B^ (salmon) with respect to the SeThsA SIR2^B^ (yellow) is indicated by black arrows. (d) Surface representation of ThsA SIR2 domains. (e) SeThsA SLOG dimer. (f) Structural superposition of SeThsA (cyan) and BcThsA (slate) SLOG dimers. Movements of BcThsA SLOG domains with respect to the SeThsA SLOG domains are indicated by black arrows. (g) Predicted model of SeThsA after 3’cADPR binding. Broken arrows indicate SeThsA SIR2 domain movements upon v2-cADPR (3’cADPR) binding to the SLOG domains. (h) NADase activities of 50 μM SeThsA mutants +/- 50 μM 3’cADPR. Initial NAD^+^ concentration was 500 μM.

### v2-cADPR (3’cADPR) production is associated with immunity suppression of the bacterial effector HopAM1

v2-cADPR (3’cADPR) is also produced by the *Pseudomonas syringae* DC3000 TIR-domain effector HopAM1 (*33*). To examine HopAM1’s ability to suppress immunity, we generated transgenic *Arabidopsis* plants that express HopAM1 and the catalytically-null mutant HopAM1^E191A^. The transgenic plants were challenged with the immunity inducing peptide flg22 (1 mM) and the production of reactive oxygen species (ROS), a hallmark of plant immunity, was quantified. Plants induced to express HopAM1 with estradiol, but not HopAM1^E191A^ or the uninduced plants, had strongly suppressed ROS production (Fig. S11b-c). The immunity suppression of HopAM1 is clearly co-related with v2-cADPR (3’cADPR) production, but not NAD^+^ depletion (Fig. S11a-d). These results indicate that v2-cADPR (3’cADPR) is responsible for HopAM1’s suppression of plant immunity.

## Discussion

Bacterial and plant TIR domains produce cyclic signaling nucleotides with immune and virulence functions, using NAD^+^ or nucleic acids as substrate (*26, 27, 33, 34, 41, 46*). Here, we report the chemical structures of two TIR domain-produced cADPR isomers, v-cADPR and v2-cADPR, which reveal that TIR domains can catalyze *O*-glycosidic bond formation between the ribose sugars in ADPR and that cyclization occurs at the 2’ (v-cADPR; 2’cADPR) and 3’ (v2-cADPR; 3’cADPR) positions of the adenosine ribose. These linkages were unexpected, because canonical cADPR produced by glycohydrolases such as CD38, *Aplysia californica* ADP-ribosyl cyclase and the SARM1 TIR domain is cyclized via the N1 position of the adenine ring (*28, 30, 47, 48*) and it had therefore been previously proposed that the cADPR isomers are likely cyclized via the alternative N positions (N6 and N7) of the adenine ring (*27, 33*). NAD^+^-dependent *O*-glycosidic bond formation between ribose sugars is a new enzymatic activity of TIR domains, but it has previously been reported for the ART (ADP-ribosyl transferase) domain of poly(ADP-ribose) polymerases (PARPs), which catalyze 2′–1″ and 2″–1″ ribose-ribose bonds between ADPR molecules using NAD^+^ as a substrate (*49, 50*).

Self-association is critical for the NADase activity of SARM1 and plant TIR domains (*25, 26, 38*) and our biochemical studies with AbTir suggest that this is also the case for cADPR isomer-producing bacterial TIR domains. In the SARM1 TIR domain, self-association is facilitated by the SAM domains, which form an octameric ring structure, while in plant TNLs, self-association of TIR domains requires the central NB-ARC domains, which form a tetrameric structure. The CC domains of both AbTIR and TcpB (*11*) self-associate in solution and may therefore have a similar role to the SARM1 SAM domains, plant TNL NB-ARC domains and bacterial SAVED domains (*51*) in facilitating TIR domain clustering.

The active-site region and self-association interfaces of SARM1 are conserved in the plant immune receptors RPP1 and ROQ1 (*6, 7, 25, 38–40*), and we recently classified the corresponding assemblies as “enzyme” TIR assemblies (*6*). They are different from the assemblies formed by the TIR domains from the TLR adaptors MAL and MyD88 (scaffold assemblies); while they both display a BB-loop-mediated head-to-tail arrangement of TIR domains, two such TIR-domain strands associate in an anti-parallel manner in enzyme TIR assemblies, while they associate in a parallel manner in scaffold TIR-domain assemblies, such as MAL and MyD88. Crystal structures of cADPR isomer-producing bacterial TIR domains AbTir^TIR^, BtTir^TIR^, TcpB^TIR^ (*11, 42, 52*) and BcThsB (*35*) do not display interfaces analogous to any of these assemblies. Different oligomeric arrangements underpin the NADase and 2′,3′-cAMP/cGMP synthetase activities of plant TIR domains (*41*). Unexpectedly, our cryo-EM structure of the filamentous assembly of AbTir^TIR^ in the presence of the NAD^+^ mimic **3AD** reveals that it adopts a scaffold assembly arrangement. Our mutagenesis data confirms that the observed arrangement is important for its catalytic function, and that an intact BB loop is also required. The active site is remarkably similar to the one in SARM1 (*38*), consistent with it being formed though the analogous BE-interface mediated association. Comparison of the monomeric AbTir^TIR^ structure to the filamentous assembly reveals remarkable conformational changes. As the additional domains in bacterial TIR proteins play a role in self-association, it will be of interest to find out how they facilitate the active configuration and what size active complexes form in bacterial cells. The symmetric interface - found in crystal structures of all bacterial TIR domain-containing proteins with known structure, except for BcThsB (*11, 12, 35, 42, 43*), and shown to be important for the ability of TcpB to self-associate and modulate Toll-like receptor signaling (*11, 53*) - may play a regulatory role in the transition to the active state.

Our studies reveal that a highly conserved tryptophan residue in the αC helical region, part of the previously defined WxxxE motif (*54*), plays a crucial role in the cyclization of ADPR by bacterial TIR domains. The equivalent tryptophan also plays an important role in SARM1 and plant TIR domains. In SARM1, this tryptophan W638 mediates aromatic stacking interactions with NAD^+^ mimetics and the W638A mutant has lower NADase activity compared to the wild-type protein (*25, 38*). In the flax L6 TNL protein, mutation of this tryptophan (W131) to an alanine abrogates cell-death signaling, and in the cryoEM structure of the related flax L7 TIR domain in complex with DNA, it is involved in interaction with the product 2′,3′-cAMP (*41, 55*). In the *Aplysia californica* ADP ribose cyclase, which is evolutionarily related to the CD38 glycohydrolase, and similar to AbTir and AaTir in that it produces a cyclic ADPR (canonical cADPR) as the major product of NAD^+^ hydrolysis, a phenylalanine residue (F174) directs the folding of the substrate during the cyclization reaction, by interacting with the adenine base of ADPR after Nam cleavage (*56, 57*). Cyclization of 1″-2′ and 1″-3′ ribose *O*-glycosidic bonds in ADPR by TIR domains will require the two ribose sugars to come into close proximity after Nam cleavage and a possible role for the conserved αC helix tryptophan residue is to facilitate such substrate folding via interaction with the adenine base.

Although multiple plantTNL proteins and bacterial TIR domain-containing proteins produce cADPR isomers, their mechanism of action and targets are only starting to be resolved. A cADPR isomer produced by ThsB TIR domains of Thoeris defence systems, but not the canonical cADPR, has recently been found to act as a second messenger upon phage infection and cause abortive infection, by activating the NADase activity of ThsA via binding to its SLOG domain (*34*). The identity of this ThsB-produced cADPR isomer was not reported; we speculate that it corresponds to v2-cADPR (3’cADPR), because our results demonstrate that this isomer is a strong activator of ThsA NADase activity and has nanomolar affinity for ThsA proteins from different bacteria. v-cADPR (2’cADPR) can also trigger the NADase activity of ThsA (Fig. 5b and (*34*)), but as it is a significantly weaker binder than v2-cADPR (3’cADPR) (Fig. 6), a much higher concentration of v-cADPR (2’cADPR) is needed to activate ThsA. Consistent with our biochemical data, the BcThsA^SLOG^:v2-cADPR (3’cADPR) complex structure reveals a conserved binding pocket that is selective for v2-cADPR (3’cADPR). Our structural data also suggest that v2-cADPR (3’cADPR) binding to the SLOG domain induces a reorganization of the ThsA tetramer to allosterically promote binding to its substrate NAD^+^. This mode of action is reminiscent of the nicotinamide mononucleotide (NMN)-induced activation of SARM1, which is only able to bind and cleave the substrate NAD^+^ after a change to its octamer organization triggered by NMN binding to its ARM domain (*38*). v-cADPR (2’cADPR) produced by the protein BdTIR from the plant *Brachipodium distachyon* has been found to bind to the protein Tad1 (Thoeris anti-defense 1) that inhibits Thoeris immunity (*58*), but there is no data showing that this is the isomer produced by ThsB proteins. It will be of interest to uncover the identity of ThsB-produced cADPR isomers and the structural basis for how they activate ThsA. SLOG domains are also found in cytokinin-activating proteins in plants (*36, 59, 60*) and it will be of interest to determine if these proteins are also receptors for v- and v2-cADPR (2’ and 3’cADPR).

The nucleotides pRib-AMP/ADP (2’-(5’’-phosphoribosyl)-5’-adenosine mono-/di-phosphate) were recently shown to trigger immune signaling in plants by allosterically promoting the EDS1 (enhanced disease susceptibility 1) - PAD4 (phytoalexin deficient 4) complex to bind to the plant NLR protein ADR1-L1 (*61*). The production of these nucleotides requires TIR domain-containing proteins, but the substrates have not been identified. Our cADPR isomer structures show that pRib-AMP can be derived directly from v-cADPR (2’cADPR) by cleavage of its pyrophosphate bond, suggesting that NAD^+^ could be the substrate. The cleavage of the pyrophosphate bond could indicate the involvement of plant NUDIX hydrolases like NUDX6/7, which regulate plant immunity by degrading 2’3’-cAMP/2’3’-cGMP, the other nucleotides putatively produced by TIR domains (*41*). Plant TIR domains can also generate ADP-ribosylated ATP (ADPr-ATP) and di-ADPR, which in turn promote the association of EDS1 and SAG101 (senescence-associated gene 101) with the helper NLR NRG1A (N requirement gene 1A) (*46*).

Interestingly, 3′-O-β-D-ribofuranosyladenosine, which has an identical 1’’-3’ *O*-glycosidic linkage to v2-cADPR (3’cADPR) but lacks the phosphate groups, has been shown to accumulate in leaves infected with the HopAM1 producing bacterium *Pseudomonas syringae* DC3000 (*62*), suggesting that cADPR isomers can be further modified in plants. Pyrophosphate bond cleavage of HopAM1 produced v2-cADPR (3’cADPR) followed by removal of the two ribose-5-phosphate groups is a possible synthetic path for this nucleotide product, which suggests that cADPR isomers perhaps not only serve as signaling molecules but are also important intermediates in the synthesis of additional novel nucleosides associated with plant immunity. In conclusion, our study unravels the cyclization site of cADPR isomers and provides new insights into nucleotide production and signaling by TIR domains.

## Acknowledgments

We acknowledge use of the Australian Synchrotron MX facility and thank the staff for their support. We acknowledge the Centre for Microscopy and Microanalysis, University of Queensland and staff (Lou Brillault, Matthias Floetenmeyer, Richard Webb and Roger Wepf). We thank Fumiaki Makino (JEOL) for help with cryo-EM data collection, and Kasun Arachchige and Jack Clegg for help with crystallization.

## Funding

The work was supported by the National Health and Medical Research Council (NHMRC grants 1196590 to T.V., 1107804 and 1160570 to B.K., and T.V., 1071659 to B.K., and 1108859 to T.V.), the Australian Research Council (ARC) Future Fellowship (FT200100572) to T.V., the ARC Laureate Fellowship (FL180100109) to B.K., the ARC DECRA (DE170100783) to T.V., the National Institutes of Health (R01NS087632 to J.M. and A.D.), Nebraska Soybean Board project No. 1734 (to M.Guo.) and Biotechnology and Biological Sciences Research Council (BB/V00400X/1) to M.Grant and L.S. Y.S. was a recipient of Griffith University Postdoctoral Fellowship Scheme.

## Author contributions

Conceptualization, M.K.M, Y.S., M.A.Z., M.Grant., M.Guo., J.M., A.D., T.V., B.K.; Investigation, M.K.M, Y.S., S.L, M.A.Z., N.D., S.E., T.G.S. W.G., V.M., T.M., J.S.W., S.J.H., E.V., L.H-T., N.M., B.Y.J.L., H.B., M.J.L., I.P., L.S., M.Gr., J.D.N., M.Guo., T.V.; Writing – original draft, M.K.M, Y.S., S.L, T.V., B.K.; Writing – review & editing, all authors; Funding acquisition, T.V., B.K., M.Gu., M.Grant, L.S., J.M., A.D.; Resources, T.V. B.K., J.M., A.D.; Supervision, T.V., B.K., J.D.N., J.M., A.D.. M.A.S., M.Grant., M.Guo.

## Competing interests

The authors have no competing interests.

## Data and materials availability

Coordinates and structure factors for the AbTir^TIR^, BtTir^TIR^, BcThsA^SLOG^:v2-cADPR (3’cADPR) and SeThsA crystal structures have been deposited in the Protein Data Bank with IDs 7UWG, 7UXR, 7UXS and 7UXT, respectively. Maps and coordinates for the AbTir^TIR^:3AD cryoEM structure have been deposited to the Electron Microscopy Data Bank (EMD-26862) and Protein Data Bank (7UXU), respectively.

## Supplementary Materials for

### Materials and Methods

#### Cloning

AbTir: ligation-independent cloning (LIC) (*63*) was used to generate the AbTIR constructs. Full-length AbTir cDNA (GenBank: EXB04249.1) was obtained as a gBlock (Integrated DNA Technologies). AbTir^full-length^ (amino acid 1-269), AbTir^CC^ (amino acid 27-118) and AbTir^TIR^ (amino acid 134-267) were amplified using AccuPower® Pfu PCR PreMix (Bioneer Pacific). Crystallization Construct Designer (https://ccd.rhpc.nki.nl/) was used to design all primers (*64*). The forward and reverse primers had the following overhangs, respectively: Fw: TACTTCCAATCCAATGCG; Rv: TTATCCACTTCCAATGTTA. The amplified products and SSpI (NEB Cat # R0132S)-digested pMCSG7 plasmids (*65*) were digested with T4 DNA polymerase (NEB). Subsequently, 2 µl of T4 DNA polymerase-treated PCR product, and pMCSG7 were incubated at room temperature for 30 minutes. The mixtures were then transformed into *E. coli* (DH5α) competent cells using a lysogeny broth (LB) agar plate containing 100 µg/mL ampicillin. The LB plate was then incubated for 16 hours at 37°C. *E. coli* colonies having the plasmids were confirmed by colony PCR using AccuPower® Taq PCR Premix (Bioneer Pacific). Four successfully transformed colonies were then grown in 10 mL LB containing 100 µg/mL ampicillin (Sigma-Aldrich) in a 50 mL Falcon tube for 16 h at 37°C. Plasmids were extracted from the cultures using QIAprep® Spin Miniprep Kit from Qiagen. Then, all constructs were sequenced using the AGRF (Australian Genome Research Centre) Sanger sequencing service.

ThsA proteins: full-length BcThsA (WP_002078322.1), BcThsA^SIR2^ (residues 1-284), BcThsA^SLOG^ (284-476) AbThsA (WP_032061149), EfThsA (WP_230207162), SeThsA (WP_012679271), and SeThsA^SIR2^ (residues 1-283) were synthesized (gBlock, Integrated DNA Technologies) and cloned into the pMCSG7 vector using LIC (*65*).

AbTir, AaTir, BtTir, BXY39700, Btheta7330_RS03065, Bovatus_RS22005, AMN69_RS28245, ORFOR_RS09155, PROVRUST_05034, AMN69_RS06490, CLOBOL_01188 for HPLC assays: DNA fragments encoding TIR domains codon-optimized for *E. coli* expression were synthesized and cloned into the pET30a(+) vector, in between NheI and HindIII restriction sites, with an N-terminal tandem Strep-tag and a C-terminal 6x-histidine tag.

#### Site-directed mutagenesis

All AbTir^TIR^ mutants were prepared by using a pair of complementary primers with the desired mutation, and AbTir^TIR^ (amino acid 134-267) in the pMCSG7 vector was used as the template. The plasmid DNA with the desired mutation was amplified using AccuPower® Pfu PCR PreMix from Bioneer Pacific. The amplified PCR products were then purified using the QIAquick PCR Purification Kit (Qiagen). The purified PCR products were then treated with DpnI (NEB Cat # R0176S) to destroy the template DNA. After DpnI digestion, *E. coli* (DH5α) competent cells were transformed with the plasmid DNA. All the colonies were screened, and the purified plasmids were sequenced using the same method as described in the cloning section.

EfThsA mutants were produced using Q5^®^ Site-Directed Mutagenesis (New England BioLabs), while SeThsA mutants were synthesized (gBlock, Integrated DNA Technologies) and cloned into the pMCSG7 vector using LIC (*65*). Pure plasmids were prepared using the QIAprep Spin Miniprep Kit (Qiagen) and the sequences confirmed by the Australian Genome Research Facility.

#### Protein expression

AbTir, AbTir^TIR^ and AbTir^CC^: for protein expression, BL21-Gold (DE3) Competent Cells (Agilent Technologies, Inc.) were transformed using the desired plasmid and grown on a LB-ampicillin (100 µg/mL) plate. The next day, 10 mL starter culture was grown for 16 hours at 37°C in LB media containing 100 µg/mL ampicillin. The following day, 1 mL of the 16-hour culture was added to 1 L autoclaved LB-ampicillin (100 µg/mL) media in 2.5 L ultra-yield flasks (Thomson’s Ultra Yield Flasks™, Genesearch). The flasks were incubated at 37°C in a shaking incubator (New Brunswick™ Innova® 44) at 225 rpm, until OD_600_ reached 0.6-0.8. After that, IPTG (isopropyl β- d-1-thiogalactopyranoside) (Merck Millipore) was added to a final concentration of 1 mM and the cultures incubated for 12-16 hours at 15°C.

AaTir^TIR^, BtTir^TIR^ and ThsA proteins: AaTir^TIR^ (residues 2-144, WP_091411838), BtTir^TIR^ (residues 156-287, WP_048697596) in the pET30a vector (N-terminal tandem Strep-tag and C-terminal His_6_-tag), and BcThsA, BcThsA^SIR2^, BcThsA^SLOG^ AbThsA, EfThsA, SeThsA and SeThsA^SIR2^ in the pMCSG7 vector (N-terminal His_6_-tag, TEV-protease cleavage site) were produced in *E. coli* BL21 (DE3) cells, using the autoinduction method (*66*) and purified to homogeneity, using a combination of immobilized metal-ion affinity chromatography (IMAC) and size-exclusion chromatography (SEC). The cells were grown at 37°C, until an OD^600^ of 0.6- 0.8 was reached. The temperature was then reduced to 20°C, and the cells were grown overnight for approximately 16 h. The cells were harvested by centrifugation at 5000 x *g* at 4°C for 15 min and stored at −80°C until used for purification.

#### Protein purification

AbTir, AbTir^TIR^ and AbTir^CC^: Cells were harvested by centrifuging at 4000 rpm (Beckman Coulter J-26 XPI, JLA 9.1 rotor) for 20 min at 4°C. After centrifugation, the supernatant was discarded, and the cell pellet was resuspended in ice-cold lysis/wash buffer (3 mL/L) (2X PBS, 300 mM NaCl, 30 mM imidazole, 1 mM phenylmethanesulfonylfluoride (PMSF)). Bacterial cell lysis was performed by using sonication (Branson, 10 seconds pulse, 10 seconds off at 40% amplitude). Lysed samples were then centrifuged (Beckman Coulter J-26 XPI, JA 20 rotor) for 40 minutes at 4°C to remove the cell debris, and the supernatant was loaded onto a 5 mL HisTrap column (GE Healthcare) at 4 mL/min. After that, the column was washed using 20 column volumes (CVs) of ice-cold lysis/wash buffer (3 mL/L) (2X PBS, 300 mM NaCl, 30 mM imidazole, 1 mM phenylmethanesulfonylfluoride (PMSF)). The protein was eluted using 10 CVs of elution buffer (100 mM Hepes pH 8.0, 500 mM NaCl, 500 mM imidazole). The eluted samples were then analyzed by 15% SDS-PAGE, and fractions containing pure proteins were pooled and dialyzed for 30 minutes in dialysis buffer (2X PBS, 1 mM DTT) at 4°C, to remove imidazole. After 30 minutes, tobacco etch virus (TEV) protease was added and incubated overnight at 4°C to remove the His-tag. The next day, the dialyzed samples were passed through a 5 mL HisTrap column (GE Healthcare) to remove the TEV protease. Then, the sample was further purified using size-exclusion chromatography (SEC) using the S75 HiLoad 26/600 column (GE Healthcare), pre-equilibrated with the gel-filtration buffer (10 mM HEPES pH 8.0, 150 mM NaCl). SEC was performed using ÄKTAprime or ÄKTA pure (GE Healthcare) systems.

AaTir^TIR^, BtTir^TIR^, BcThsA, AbThsA, EfThsA and SeThsA: The cells were harvested by centrifugation at 5000 x *g* at 4°C for 15 min, the cell pellets were resuspended in 2-3 mL of lysis buffer (50 mM HEPES pH 8.0, 500 mM NaCl) per g of cells. The resuspended cells were lysed using a digital sonicator and clarified by centrifugation (15,000 x *g* for 30 minutes). The clarified lysate was supplemented with imidazole (final concentration of 30 mM) and then applied to a nickel HisTrap column (Cytiva) pre-equilibrated with 10 CVs of the wash buffer (50 mM HEPES pH 8.0, 500 mM NaCl, 30 mM imidazole) at a rate of 4 mL/min. The column was washed with 10 CVs of the wash buffer followed by elution of bound proteins using elution buffer (50 mM HEPES pH 8, 500 mM NaCl, 250 mM imidazole). The elution fractions were analysed by SDS-PAGE and the fractions containing the protein of interest were pooled and further purified on either a S75 HiLoad 26/600 column (AaTir^TIR^ and BtTir^TIR^) or a S200 HiLoad 26/600 column (BcThsA, AbThsA, EfThsA and SeThsA) pre-equilibrated with gel-filtration buffer. The peak fractions were analysed by SDS-PAGE, and the fractions containing AaTir, BtTir or ThsA were pooled and concentrated to final concentrations of approximately 11.2 mg/mL (AaTir^TIR^), 4.1 mg/mL (BtTir^TIR^), 34.4 mg/mL (BcThsA), 46.2 mg/mL (AbThsA), 37.1 mg/mL (EfThsA) and 39.5 mg/mL (SeThsA), flash-frozen as 10 µL aliquots in liquid nitrogen, and stored at −80°C.

BcThsA^SLOG^, BcThsA^SIR2^ and SeThsA^SIR2^: The cells were harvested by centrifugation at 5000 x *g* at 4°C for 15 min, the cell pellets were resuspended in 2-3 mL of lysis buffer (50 mM HEPES pH 8.0, 500 mM NaCl) per g of cells. The resuspended cells were lysed using a digital sonicator and clarified by centrifugation (15,000 x *g* for 30 minutes). The clarified lysate was supplemented with imidazole (final concentration of 30 mM) and then applied to a nickel HisTrap column (Cytiva) pre-equilibrated with 10 CVs of the wash buffer (50 mM HEPES pH 8.0, 500 mM NaCl, 30 mM imidazole) at a rate of 4 mL/min. The column was washed with 10 CVs of the wash buffer, followed by elution of bound proteins using elution buffer (50 mM HEPES pH 8, 500 mM NaCl, 250 mM imidazole). The elution fractions were analysed by SDS-PAGE and the fractions containing BcThsA^SLOG^, BcThsA^SIR2^ or SeThsA^SIR2^ were pooled, supplemented with TEV protease and dialysed into gel-filtration buffer (10 mM HEPES pH 7.5, 150 mM NaCl) for 16-20 h. After dialysis, cleaved BcThsA^SLOG^, BcThsA^SIR2^ or SeThsA^SIR2^ was reloaded onto the HisTrap column to remove the TEV protease, His_6_-tag and contaminants. After the second IMAC step, BcThsA^SLOG^, BcThsA^SIR2^ or SeThsA^SIR2^ were further purified a S200 HiLoad 26/600 column pre-equilibrated with gel-filtration buffer. The peak fractions were analysed by SDS-PAGE, and the fractions containing BcThsA^SIR2^ or SeThsA^SIR2^ were pooled and concentrated to final concentrations of approximately 49 mg/ml (BcThsA^SLOG^), 8.3 mg/mL (BcThsA^SIR2^), and 32 mg/mL (SeThsA^SIR2^), flash-frozen as 10 µL aliquots in liquid nitrogen, and stored at −80°C.

AbTir, BXY39700, Btheta7330_RS03065, Bovatus_RS22005, AMN69_RS28245, ORFOR_RS09155, PROVRUST_05034, AMN69_RS06490, CLOBOL_01188 for HPLC assays: expression vectors were transformed into *E. coli* (NEB Iq/LysY, catalog number 3013I). Single colonies were grown overnight in LB with kanamycin, diluted 100x in LB, and shaken at 30°C to mid-exponential phase (OD 0.4-0.8). Protein expression was induced by adding IPTG to a final concentration of 0.1 mM and shaking at 30°C for 3 hours. Cultures were pelleted by centrifugation then resuspended in binding buffer (100 mM Tris HCl, 150 mM NaCl, pH 8.0). 10x protease inhibitor cocktail was added, samples were lysed by sonication, and lysates were clarified by ultracentrifugation. 200 µL of streptactin magnetic bead suspension (PureCube-HiCap Streptactin MagBeads, Cube Biotech), washed three times with binding buffer, was suspended with each lysate sample and incubated for 1 h at 4°C with gentle agitation. Protein-laden beads were washed three times with binding buffer and resuspended in 200 µL of binding buffer.

#### Fluorescence-based NADase assay

1, N^6^-ethenoNAD (εNAD) (Sigma-Aldrich), a fluorescent analog of NAD^+^, was used as the substrate in this assay (*25, 67*). The assay was carried out in 96-well microplate (Greiner). Fluorescence intensity was measured using a CLARIOstar® microplate reader (excitation wavelength 310-330 nm; emission wavelength 390-410 nm; readings every 1-3 minutes over 4 hours at 25 °C). The change in fluorescence over time was calculated from the slopes of the linear component of the curves. For all the fluorescence-based NADase assays, 100 µM protein and 100 µM substrate were used. The data was analyzed by Microsoft Excel and Prism GraphPad.

#### Thin layer chromatography (TLC)

TLC was performed on 5 × 10 cm silica gel 60 F_254_ plates (Merck). For TLC, 1 mM ligand was incubated with 100 μM protein on ice for 12 hours. The samples (2 μL) were then spotted into the plate and separation was performed in n-propanol/ammonium hydroxide/water (13:6:1). The plate was air-dried, and bands were visualized with a short-wavelength (254 nm) ultraviolet light source.

#### NAD^+^-cleavage product quantification by HPLC

In vitro reactions consisted of 10 µL of protein-laden bead suspension and 40 µL of 10 µM NAD^+^ in 25 mM HEPES buffer (pH 7.5) at room temperature with constant agitation. Reactions were quenched at 1 h or 48 h by pulling the beads to the side and transferring 40 µL of the reaction mixture to a new tube containing 160 µL of ice-cold 0.5 M HClO_4_. Acid metabolite extracts were spun at 20,400 x *g* for 10 minutes at 4°C. 150 µL of supernatant were neutralized with 16 µL of 3 M K_2_CO_3_ and again spun at 20,400 x *g* for 10 min at 4°C. 90 µL of supernatant were mixed with 10 µL 0.5 M potassium phosphate buffer. Metabolites were analyzed by HPLC (Shimadzu LC40) using a C18 analytical column (Kinetex, 100 x 3 mm, Phenomenex).

#### NMR-based NADase assay

NMR samples were prepared in 175 µL HBS buffer (50 mM HEPES, 150 mM NaCl, pH 7.5), 20 µL D_2_O, and 5 µL DMSO-d6, resulting in a total volume of 200 μL. Each sample was subsequently transferred to a 3 mm Bruker NMR tube rated for 600 MHz data acquisition. All ^1^H NMR spectra were acquired with a Bruker Avance 600 MHz NMR spectrometer equipped with ^1^H/^13^C/^15^N triple resonance cryoprobe at 298 K. To suppress resonance from H_2_O, a water-suppression pulse program (P3919GP), using a 3-9-19 pulse-sequence with gradients (*68, 69*), was implemented to acquire spectra with an acquisition delay of 2 s and 32 scans per sample. For each reaction, spectra were recorded at 10 min, 2 h, 4 h, 8 h, 16 h, 40 h, and 64 h time-points, depending on instrument availability. All spectra were processed by TopSpin™ (Bruker) and Mnova 11 (Mestrelab Research). The amount of NAD^+^ consumption was calculated based on the integration of non-overlapping resonance peaks, which vary depending on sample composition, from NAD^+^ and Nam, respectively. The detection limit (signal-to-noise ratio > 2) was estimated to be 10 µM.

#### STD-NMR

Samples for STD-NMR were prepared in similar solutions as for NMR NADase asasys. With a total volume of 200 µL, each sample consisted of 175 µL HBS buffer, 20 µL D_2_O, and 5 µL DMSO-d6. STD-NMR spectra were acquired with the same NMR spectrometer as for the NADase assays. The pulse-sequence STDDIFFGP19.3, in-built within the TopSpin^TM^ program (Bruker), was employed to acquire STD-NMR spectra (*70*). This pulse-sequence consists of a 3-9-19 water-suppression pulse, the parameters of which were obtained from the water-suppression pulse program (P3919GP), to suppress the resonance from H_2_O. The on-resonance irradiation was set close to protein resonances at 0.8 ppm, whereas the off-resonance irradiation was set far away from any protein or ligand resonances at 300 ppm. A relaxation delay of 4 s was used, out of which a saturation time of 3 s was used to irradiate the protein with a train of 50 ms Gaussian shaped pulses. The number of scans was 512. All spectra were processed by TopSpin™ (Bruker) and Mnova 11 (Mestrelab Research).

#### Production and purification of v-cADPR and v2-cADPR

Production reactions for v-cADPR and v2-cADPR were performed using conditions similar to the ^1^H NMR NADase assays. Each reaction was carried out in HBS buffer (50 mM HEPES, 150 mM NaCl, pH 7.5). For v-cADPR production, 1 μM of His_6_-tagged AbTir, and 10 mM NAD^+^ were added to the mixture. For v2-cADPR production, 10 μM of His_6_-tagged AaTir, and 20 mM NAD^+^ were added to the mixture. All reactions were performed at room temperature and monitored intermittently by ^1^H NMR. To stop the reaction, the His_6_-tagged protein was removed by incubating the mixture with 200 mL of HisPur™ Ni-NTA resin for 30-60 min. The resin was subsequently removed by centrifugation at 500 x *g* for 1 min and the supernatant was subjected to HPLC-based separation to purify the products. A Shimadzu Prominence HPLC equipped with a Synergi™ 4 µm Hydro-RP 80 Å column was used for separation. The mobile phase consisted of phase A (0.05 % (v/v) formic acid in water) and phase B (0.05 % (v/v) formic acid in methanol). Different gradients, flow rates, and run times were applied, depending on prior optimization with individual reaction mixtures. Product peaks were confirmed by comparison with individual chromatograms of NAD^+^, Nam and ADPR. Fractions corresponding to the product peaks were collected, concentrated, and lyophilized and stored at −20°C. For v2-cADPR production by HopAM1 in plants, HopAM1 was transiently expressed in *N. benthamiana* leaves in an estradiol-inducible plant binary vector. *N. benthamiana* leaves were ground with a mortar and pestle in liquid nitrogen. The ground powders were resuspended in 10 mL of 50% methanol kept at −40 °C and then mixed with 10 mL chloroform at −40°C. Samples were then centrifuged at 15,000 x *g* for 10 min at 0 °C and the aqueous/methanol layer was removed. The extract was lyophilized and stored at −80°C until HPLC. The v2-cADPR was purified by manual fractionation with HPLC.

#### NMR structure determination of cADPR isomers

Purified v-cADPR and v2-cADPR were used to determine their structures. At the Griffith University facility, 4 mg of v-cADPR and 4.9 mg of v2-cADPR were dissolved in 560 μL of D_2_O, respectively. Each sample was transferred to a 5 mm NMR tube rated for 600 MHz. The same NMR spectrometer as described above was utilized to acquire ^1^H, ^13^C, ^1^H -^1^H COSY, ^1^H-^13^C HSQC, and ^1^H-^13^C spectra at 298 K. The chemical structure of each compound was determined by assignments of ^1^H and ^13^C peaks and correlations, especially those linking two ribose rings (Fig. 1, Table S1–2). At the University of Warwick facility, samples were dissolved in D_2_O and ^1^H, COSY, HSQC, HMBC, NOESY spectra were acquired on Bruker Avance II 700 MHz spectrometer equipped with TCI cryoprobe. The sample was also used to acquire ^1^H-^31^P HMBC on a Bruker 600 MHz spectrometer with a BBO probe. All experiments were done at 25 °C. For HopAM1-produced v2-cADPR, HPLC-purified and lyophilized compound was reconstituted in 160 μL of deuterium oxide (D_2_O) and transferred into a 3-mm NMR tube. The samples were analyzed with a Bruker Avance-III HD 700 MHz NMR system equipped with a 5 mm QCI-P cryoprobe or a Bruker Avance NEO 600 MHz NMR system equipped with a TCI-H/F cryoprobe. The chemical structure of the compound was determined by assignment of 1- and 2-dimensional NMR data, including ^1^H, ^13^C, ^1^H-^1^H COSY, NOESY, ^1^H-^13^C HSQC, ^1^H-^13^C HMBC, ^1^H-^13^C HSQC-TOCSY, and ^1^H-^31^P HSQC-TOCSY (Table S3).

#### LC-MS/MS analysis

A Waters Xevo TQXS triple quadruple mass spectrometer coupled with Waters I-class UPLC was used for LC-MS/MS analysis of v-cADPRs from both in vitro products of AbTir and AaTir NAD^+^ activity and plant compounds extracted in 10% methanol, 1% acetic acid. The mass spectrometer is equipped with an electrospray ionization source in positive ion mode. Source condition: capillary voltage: 800V, desolvation temperature: 600°C, desolvation gas: 1000 L/h, congas 150 L/h and nebuliser gas: 7 bar. MRM transitions for v-cADPRs are parent ions at m/z 542.00 and daughter ions at 136.00 and 348.02 with collision energy at 32 and 28 eV, respectively. UPLC mobile phases comprise A; water with 2 mM ammonium acetate, and B; 100% methanol. The elution gradient was: 0-5 min, 100% A, 5-7 min, 80% A, 7-8 min, 100% B, then isocratic for 2 min at 100% B before equilibrating back to 100% A for 15 min. Flow rate was set at 0.2 mL/min. The column used was a Waters Acquity UPLC CSH C18, 1.7 μm, 0.1×100 mm. High resolution measurements were done on a Bruker MaXis II Q-TOF mass spectrometer.

#### Size-exclusion chromatography (SEC)-coupled multi-angle light scattering (MALS)

A DAWN HELEOS II 10-angle light-scattering detector coupled with an Optilab rEX refractive index detector (Wyatt Technology), combined with a Superdex 200 5/150 Increase size exclusion column (Cytiva), connected to a Prominence HPLC (Shimadzu), was used for SEC-MALS. The column was equilibrated in gel-filtration buffer, and 30 µL of the purified proteins were run through the column at 0.25 mL/min. Molecular masses were calculated using Astra 6.1 (Wyatt Technology).

#### Isothermal titration calorimetry (ITC)

ITC experiments were performed in duplicate on Nano ITC (TA Instruments). All proteins and compounds were dissolved in a buffer containing 10 mM HEPES (pH 7.5) and 150 mM NaCl. The baseline was equilibrated for 600 s before the first injection. 0.3 mM v-cADPR (2’cADPR) or v2-cADPR (3’cADPR) was titrated as 30 injections of 1.44 μL every 200 s into 50-112.4 μM AbThsA, or 20 injections of 1.44 µL every 200 s into 24-37 µM EfThsA. The heat change was recorded by injection over time and the binding isotherms were generated as a function of molar ratio of the protein solution. The dissociation constant (K_d_) values were obtained after fitting the integrated and normalized data to a single-site binding model using NanoAnalyze (TA Instruments).

#### Protein crystallization

AbTir^TIR^ crystals were obtained using the hanging-drop vapour diffusion method. Initial trays were made using the mosquito® crystallization robot (SPT Labtech). Several initial hits were obtained within a day in different commercial crystallization screens (Hampton Research Index Screen (HR2-144), Molecular Dimensions JCSG-plus Screen (MD1-37) and Molecular Dimensions SG1 Screen (MD 1-88)). Diffraction-quality crystals of AbTir^TIR^ were produced using 0.1 M Bis-Tris pH 5.5, 0.2 M LiSO_4_, and 25% PEG 3350 at 20°C. EasyXtal 15-Well Tool (Qiagen) was used for the optimization of the crystals.

BtTir^TIR^: diffraction-quality crystals were grown by the hanging drop vapour diffusion method at 293 K, with drops containing 1 μL of protein (20 mg/mL), and 1 μL of reservoir solution (0.1 M Hepes pH 7.0, 0.2 M MgCl_2_ and 16-22% PEG 3350); they appeared within a week.

BcThsA^SLOG^:3’cADPR: diffraction-quality crystals were grown by the hanging drop vapour diffusion method at 293 K, with drops containing 1 μL of protein (10 mg/mL), and 1 μL of reservoir solution (0.1 M Bis-Tris pH 5.5, 0.1 M ammonium sulfate and 25-29% PEG 3350); they appeared within a week.

SeThsA: diffraction-quality crystals were grown by the hanging drop vapour diffusion method at 293 K, with drops containing 1 μL of protein (5.5 mg/mL), and 1 μL of reservoir solution (0.1 M Mes pH 6.0, 0.2 M potassium sodium tartrate tetrahydrate and 28-30% PEG smear low (*71*)); they appeared within a week.

#### Crystallographic data collection

AbTir^TIR^: The crystals were harvested using 18 mm Mounted CryoLoop™ - 20 micron (Hampton) and cryoprotected using 50% well solution + 30% PEG 400 or 50% well solution + 30% glycerol. The harvested crystals were immediately flash-cooled in liquid nitrogen. X-ray diffraction data was collected using a wavelength of 0.9537 Å at the Australian Synchrotron MX2 beamline. Diffraction data was collected using the Blu-Ice software and indexed and integrated using XDS (*72*). Data scaling was done with Aimless in the CCP4 suite (*73*). Crystal structures were solved by molecular replacement with Phaser (*74*), using TcpB^TIR^ (PDB: 4LQC) as the search model. Model building and structure refinement were performed using Coot (*75*) and Phenix-refine (*74*), respectively. Data processing and refinement statistics are given in Table S4.

BtTir^TIR^: The crystals were cryoprotected in 20% glycerol and flash-cooled at 100 K. X-ray diffraction data were collected from single crystals on the MX2 beamline at the Australian Synchrotron, using a wavelength of 0.9537 Å. The datasets were processed using XDS (*72*) and scaled using Aimless in the CCP4 suite (*73*). The structure was solved by molecular replacement using Phaser (*76*) and the AbTir structure as a template. The models were refined using Phenix (*77*), and structure validation was performed using MolProbity (*78*). Data processing and refinement statistics are given in Table S4.

BcThsA^SLOG^:3’cADPR: The crystals were cryoprotected in 20% glycerol and flash-cooled at 100 K. X-ray diffraction data were collected from single crystals on the MX2 beamline at the Australian Synchrotron, using a wavelength of 0.9537 Å. The datasets were processed using XDS (*72*) and scaled using Aimless in the CCP4 suite (*73*). The structure was solved by molecular replacement using Phaser (*76*) and the SLOG domain of the BcThsA crystal structure (PDB: 6LHX) as a template (*35*). The models were built and refined using Phenix (*77*) and Coot, and structure validation was performed using MolProbity (*78*). Data processing and refinement statistics are given in Table S4.

SeThsA: The crystals were cryoprotected in 20% glycerol and flash-cooled at 100 K. X-ray diffraction data were collected from single crystals on the MX2 beamline at the Australian Synchrotron, using a wavelength of 0.9537 Å. The datasets were processed using Mosflm (*79*) and scaled using Aimless in the CCP4 suite (*73*). The structure was solved by molecular replacement using Phaser (*76*) and an AlphaFold2 model of SeThsA as a template (*80*). The models were built and refined using Phenix (*77*), Coot and ISOLDE (*81*) and structure validation was performed using MolProbity (*78*). Data processing and refinement statistics are given in Table S4.

#### Electron microscopy

Negative-stain electron microscopy: after dilution, AbTIR^TIR^ (at 5 mg/mL) was incubated with 2 mM **3AD** at 25°C for 1 h. Protein was diluted to 0.1 mg/mL in gel filtration buffer (containing 30 mM HEPES, pH 7.5, and 150 mM NaCl) and 2 mM **3AD,** before being loaded onto grids. 6 μL sample was placed on a carbon-coated copper gird and incubated for 5 min. The grid was then washed with gel filtration buffer containing 2 mM **3AD**, stained with 2% uranyl acetate for 30 s and air-dried. The images were collected on the Hitachi HT7700 120kV transmission electron microscope at 25,000x magnification at 120 keV.

Cryo-EM sample preparation and data collection: AbTir^TIR^ was diluted to 5 mg/mL and incubated with 2 mM **3AD** at 25°C for 1 h. Protein was then diluted to 2.5 mg/mL with gel filtration buffer containing 2 mM **3AD**, before being loaded onto grids. Quantifoil Au R 1.2/1.3 300 mesh holey carbon girds were glow-discharged for 30 s at medium level after 1 min evacuation of both carbon and copper sides. A volume of 2 μL of sample solution was added to the grids, and samples were vitrified in a Leica EMGP2 plunge freezer using a blotting time of 8.5 s at 8°C, with humidity of 96%. Screening and data collection were performed on a JEOL Cryo-ARM300 operated at 300 keV and equipped with an in column Ω energy filter (slit width 20 eV) and Gatan K3 direct detection device. Cryo-EM data collection settings are summarized in Table S5.

Data processing and 3D reconstruction: all data processing was performed with cryoSPARC (*82*) and the cryo-EM processing workflow is summarized in Figure S4. Filaments were auto-picked using filament tracer in cryoSPARC. Several rounds of 2D classification were performed to remove inferior particles. After 2D classification, good particles were further classified into three 3D maps using ab initio reconstruction. The best reconstruction was used as a reference for helical refinement. The final resolution of the 3D reconstruction is 3.41 Å.

Model building and refinement: The crystal structure of AbTir^TIR^ was docked into the electrostatic potential map in ChimeraX (*83*) and fit using ISOLDE (*81*). **3AD** and structural differences were manually built or adjusted in ISOLDE and Coot (*75*). Models were refined using multiple rounds of phenix.real_space_refine (*84*).

#### Phytobacterial challenges

Five-week old *Arabidopsis thaliana* Col-0 grown under short days (8 h light, 16 h dark, 120 microeinsteins, 65% relative humidity) were challenged with *Pseudomonas syringae* pv. tomato strain DC3000 or its 28 effector deleted derivative (D28E) OD_600_ 0.15 and left for 18 h under lights. Challenged leaves were harvested 18 h later, snap-frozen in liquid nitrogen and freeze-dried. Samples were processed for LC-MS/MS as previously described (*62*).

#### ROS assay

ROS production was determined as previously described (*85*). Briefly, transgenic *Arabidopsis* leaves were sprayed with 20 µM estradiol containing 0.02% Silwet-L77. After 24 hours, leaf discs were excised using a 4 mm diameter cork borer and incubated in H_2_O in white 96-well microtiter plates overnight. The H_2_O was then replaced with 0.5 mM luminol-based chemiluminescent probe L-012 and 1 mM flg22 in 10 mM MOPS-KOH buffer (pH 7.4). The production of ROS was determined by counting photons using with a Synergy 5 luminometer (BioTek, Winooski, VT, USA).

#### Bioinformatic analysis

To identify positions important for determining the product specificity of TIR domain NADases, we aligned 278 TIR-domain sequences using HMM. Positions where >20% of the sequences contained gaps relative to the HMM profile to which they were aligned were excluded (110 out of 116 positions were kept). Additionally, sequences that lacked significant similarity to the profile were removed (bitscore <0), as were sequences that contained many gaps relative to the model (gaps at >15% of the 110 positions). The trimmed and filtered alignment yielded 267 TIR domain sequences and 110 positions; the alignment was subsequently used to calculate the mutual information (MI) (i) between TIRs that possessed cyclase activity and those that produced ADPR, and (ii) among TIRs that produced different forms of cADPR. The position corresponding to W204 in AbTir was identified as being highly informative (MI in 90^th^ percentile) for cyclase activity, with the second lowest MI of all positions when calculating MI between TIR domains making the three different cyclic products (cADPR, v-cADPR and v2-cADPR), behind only the catalytic glutamate.

**Fig. S1.**
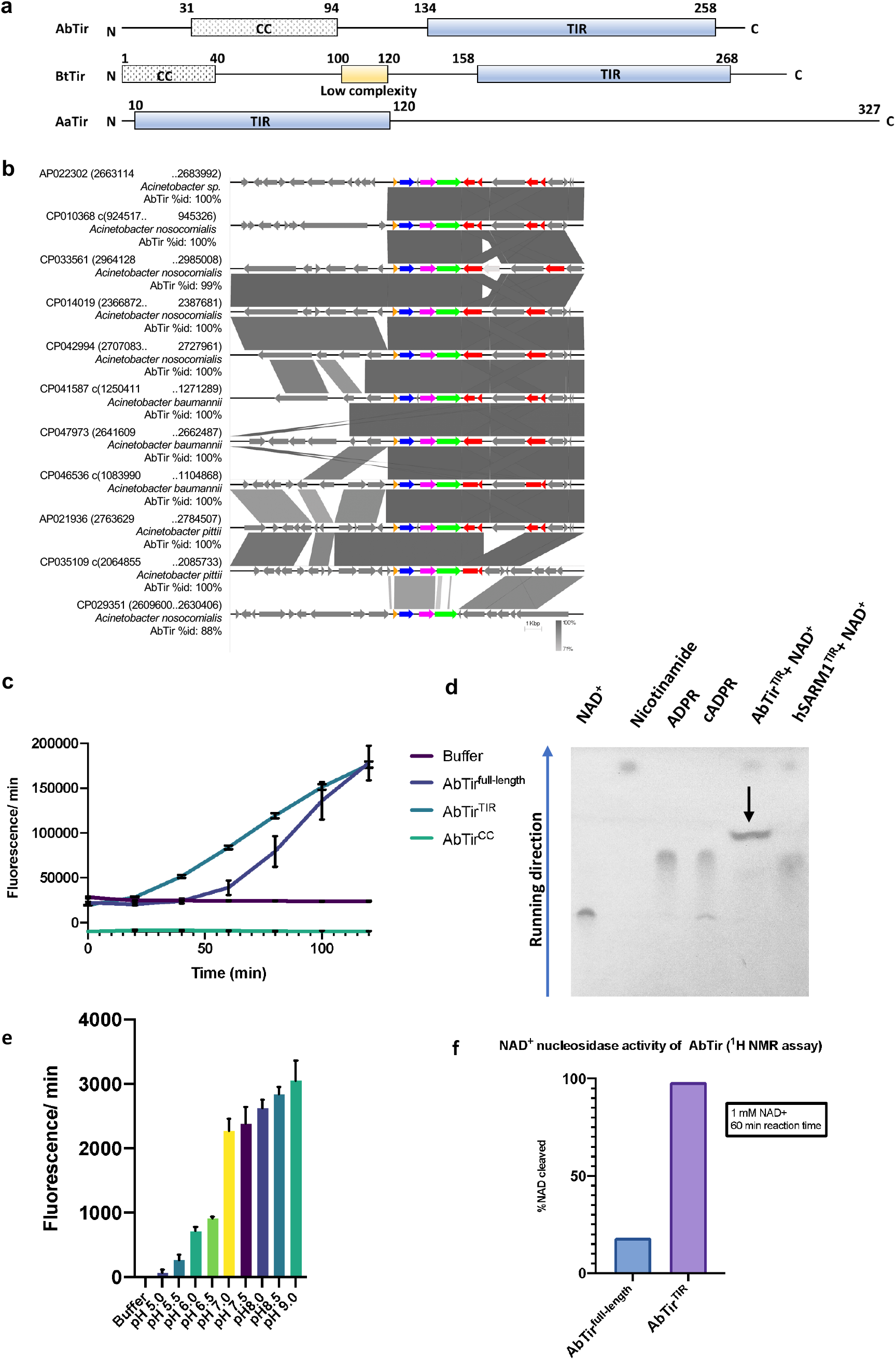
Genomic location and enzymatic characterization of AbTir^TIR^. (a) Schematic diagram of the domain organization of AbTir, BtTir, and AaTir. CC, coiled coil domain; TIR, Toll/interleukin-1 receptor (TIR) domain. (b) Pairwise sequence comparison of 10 kilobases up- and down-stream of the gene encoding AbTir. Greyscale bars represent the level of nucleotide sequence identity for that region, as indicated by the scale. Sequence annotations are colour-coded by function: blue, AbTir; orange, integrase; pink, chromate resistance; green, chromate transporter; red, IS*3* family insertion sequence; salmon, other insertion sequence; and grey, other/unknown function. We identified AbTIR in 11 complete *Acinetobacter* genomes; in each case, AbTir was located between an integrase gene and two genes encoding chromate resistance and transporter proteins, which are adjacent to 0-2 copies of an insertion sequence belonging to the IS*3* family. (c) NADase activity of AbTir^full-length^, TIR domain, CC domain and the catalytic glutamate mutant of the TIR domain. Data are presented as mean ± SD (n = 3). (d) TLC analysis of AbTir^TIR^, showing the production of v-cADPR upon cleavage of NAD^+^. (e) Impact of pH on the enzymatic activity of AbTir^TIR^. Data are presented as mean ± SD (n = 3). (f) ^1^H NMR assay for AbTir to determine NADase activity.

**Fig. S2.**
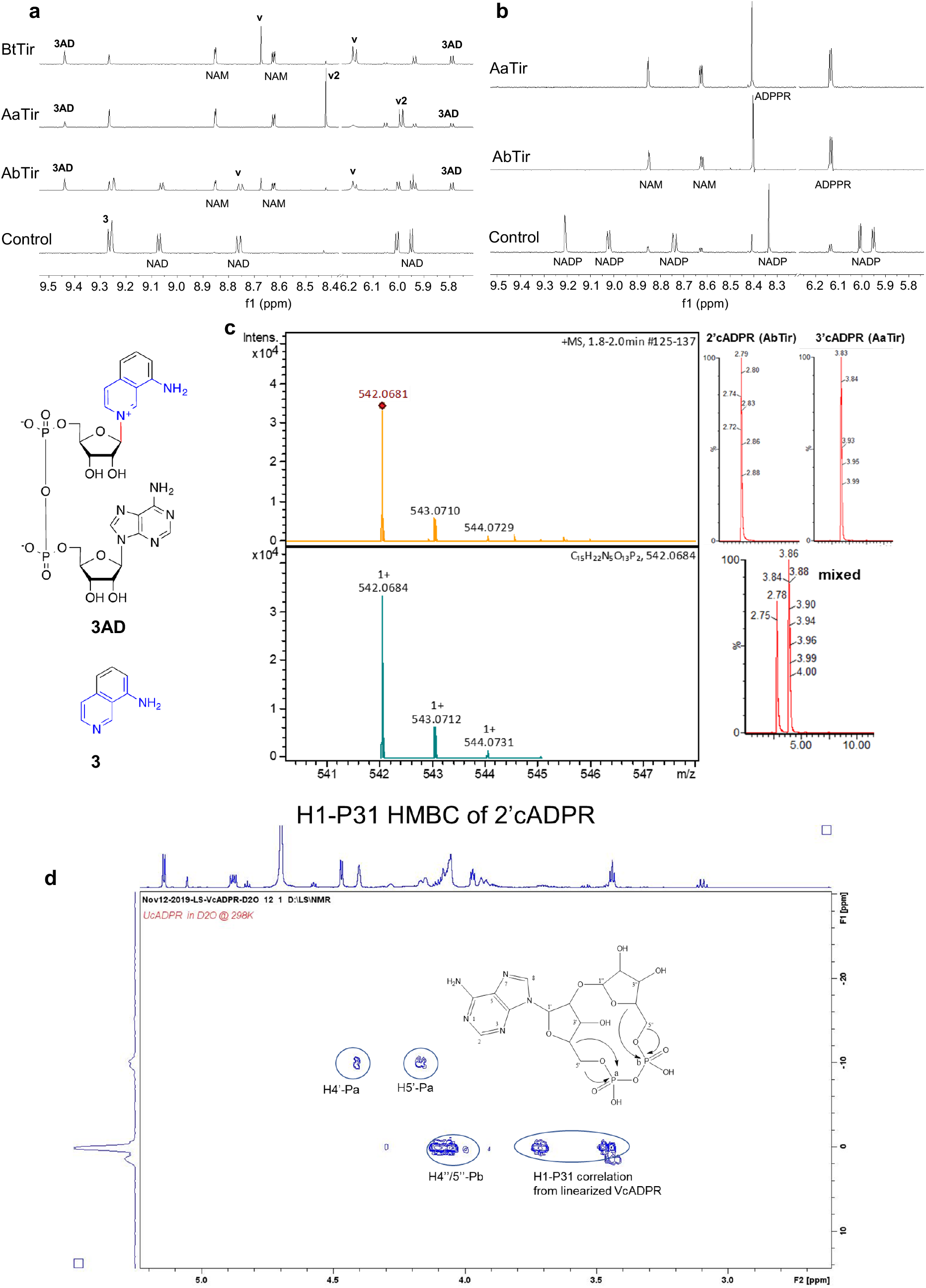
NMR and MS analyses of cADPR isomers. (a) Expansions of ^1^H NMR spectra showing base-exchange reactions by 0.1 μM AbTir, 0.5 μM AaTir, and 2.5 μM BtTir, respectively. The initial concentration for both NAD^+^ and **3** (8-amino-isoquinoline) was 500 μM. Spectra for AbTir and BtTir correspond to 40 h incubation time, while for AaTIR the incubation time was 16 h. Selected peaks are labelled, showing the formation of base-exchange product **3AD** for both proteins, as well as the production of v-cADPR (2’cADPR) (**v**) and v2-cADPR (3’cADPR) (**v2**). (b) Expansions of ^1^H NMR spectra showing hydrolysis of NADP to NAM and ADPPR by AbTir and AaTir. The initial concentration of NADP was 500 μM, while the protein concentration was 0.5 μM. All spectra correspond to 16 h incubation time. Selected peaks are labelled. (c) LC-MS/ MS of the 2’cADPR and 3’cADPR isomers produced by AbTir^TIR^ and AaTir^TIR^ NADase activity respectively. (Left) High resolution mass spectrum of AbTIR 2’cADPR. Top; measured spectrum, bottom; simulated spectrum. (Right) LC-MS/MS of AbTIR and AaTIR reveal the distinct 2’cADPR and 3’cADPR isomers. (d) ^1^H1-^31^P HMBC of v-cADPR (2’cADPR).

**Fig. S3.**
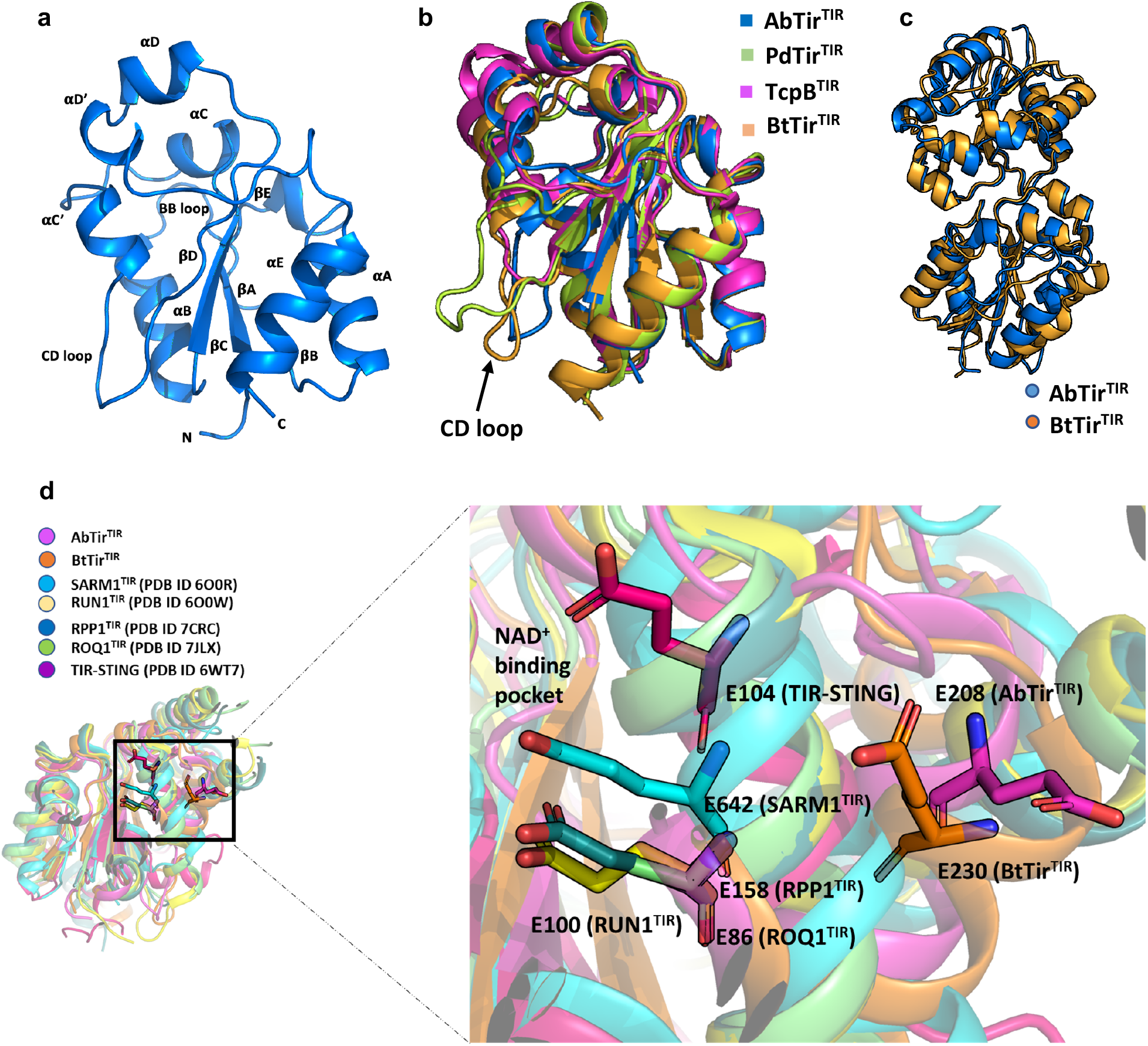
Crystal structure of AbTir^TIR^and BtTir^TIR^. (a) Crystal structure of AbTir^TIR^. (b) Structural superposition of AbTir^TIR^ and BtTir^TIR^ with PdTir^TIR^ (PDB: 3H16) and TcpB^TIR^ (PDB: 4C7M), showing the differences between the structures. (c) Structural superposition of AbTir^TIR^ and BtTir^TIR^ homodimers observed in the crystal structure, coloured blue and orange. (d) Structural superposition of the catalytic glutamate of AbTir^TIR^ with other NAD-consuming TIR domains.

**Fig. S4.**
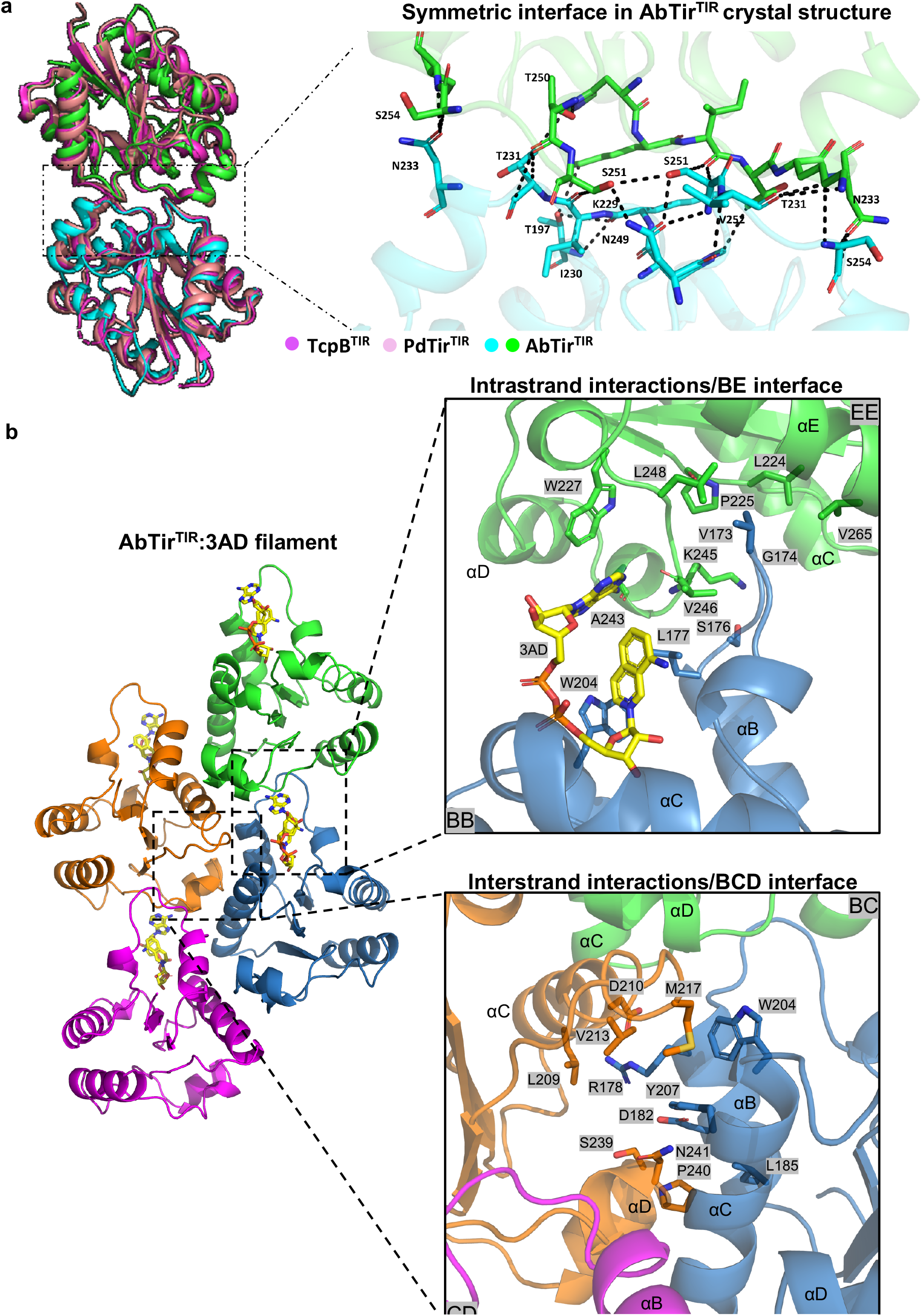
Detailed interactions within the AbTir^TIR^ crystal and cryoEM structures. (a) Left panel: Structural superposition of the symmetric dimer interface of AbTir with PdTir^TIR^ and TcpB^TIR^ (4LZP). The two molecules of AbTir^TIR^ are coloured green and cyan, respectively; the extra helix from TcpB structure is removed for better comparison and visualization. Right panel: close-up view of the interacting residues of the symmetric dimer interface of AbTir^TIR^. (b) Detailed interactions within the AbTir^TIR^:3AD filament. BB surface consist of residues in BB loop; EE surface consist of residues in βD and βE strands, and the αE helix; BC surface consist of residues in αB and αC helices; whereas CD surface consist of residues in CD loop and the αD helical region.

**Fig. S5.**
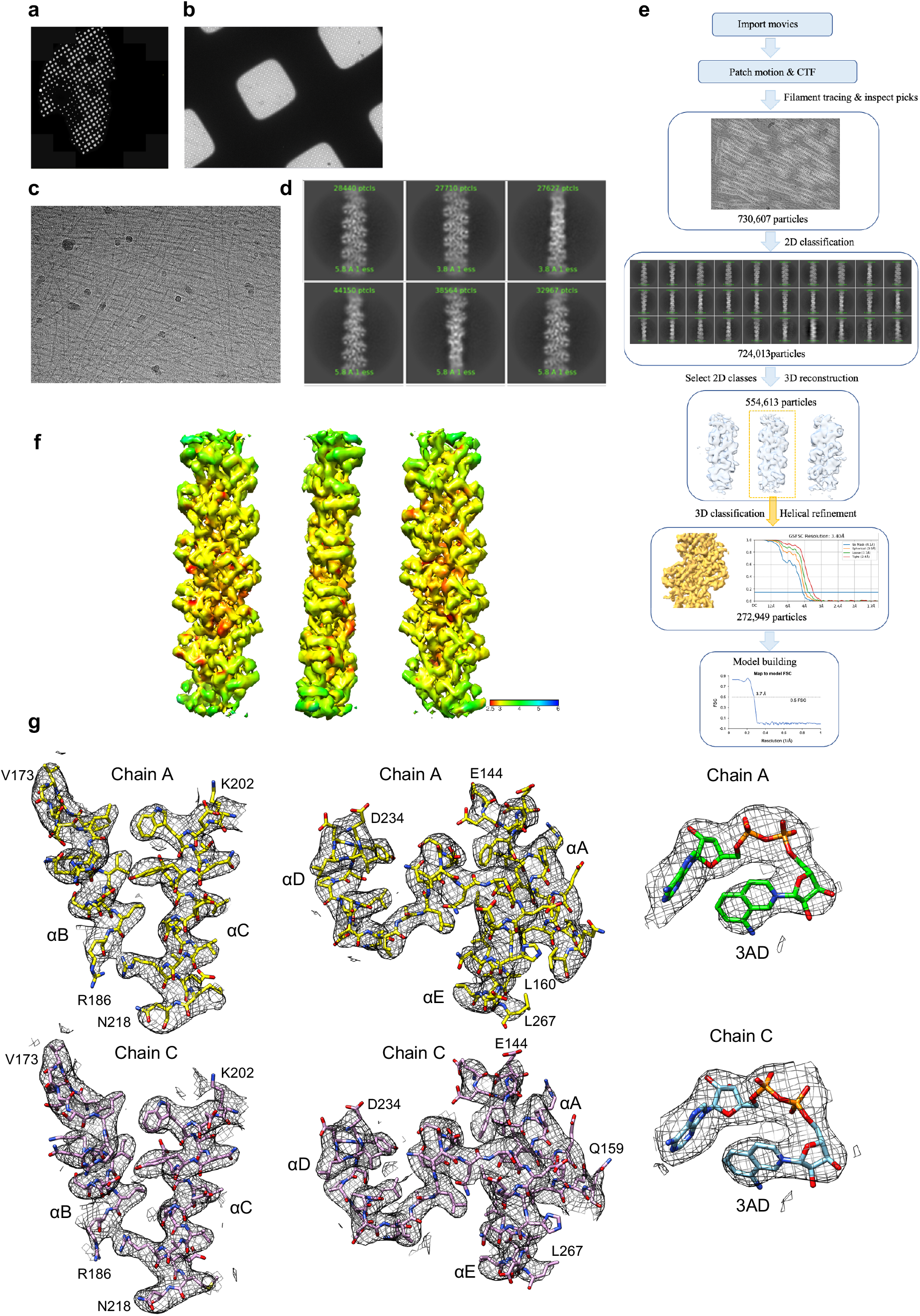
Structure determination of the AbTir^TIR^:3AD complex by cryoEM. (a-c) Representative low and high magnification cryo-EM micrographs. (d) Representative 2D class averages. (e) Flow-chart of the cryo-EM processing steps, gold-standard FSC curves of the final 3D reconstruction, and map-to-model FSC curve of the final model and the the electrostatic potential density map. (f) Local-resolution distribution of the final map. (g) Representative regions of electrostatic potential maps for 3AD-bound AbTir^TIR^.

**Fig. S6.**
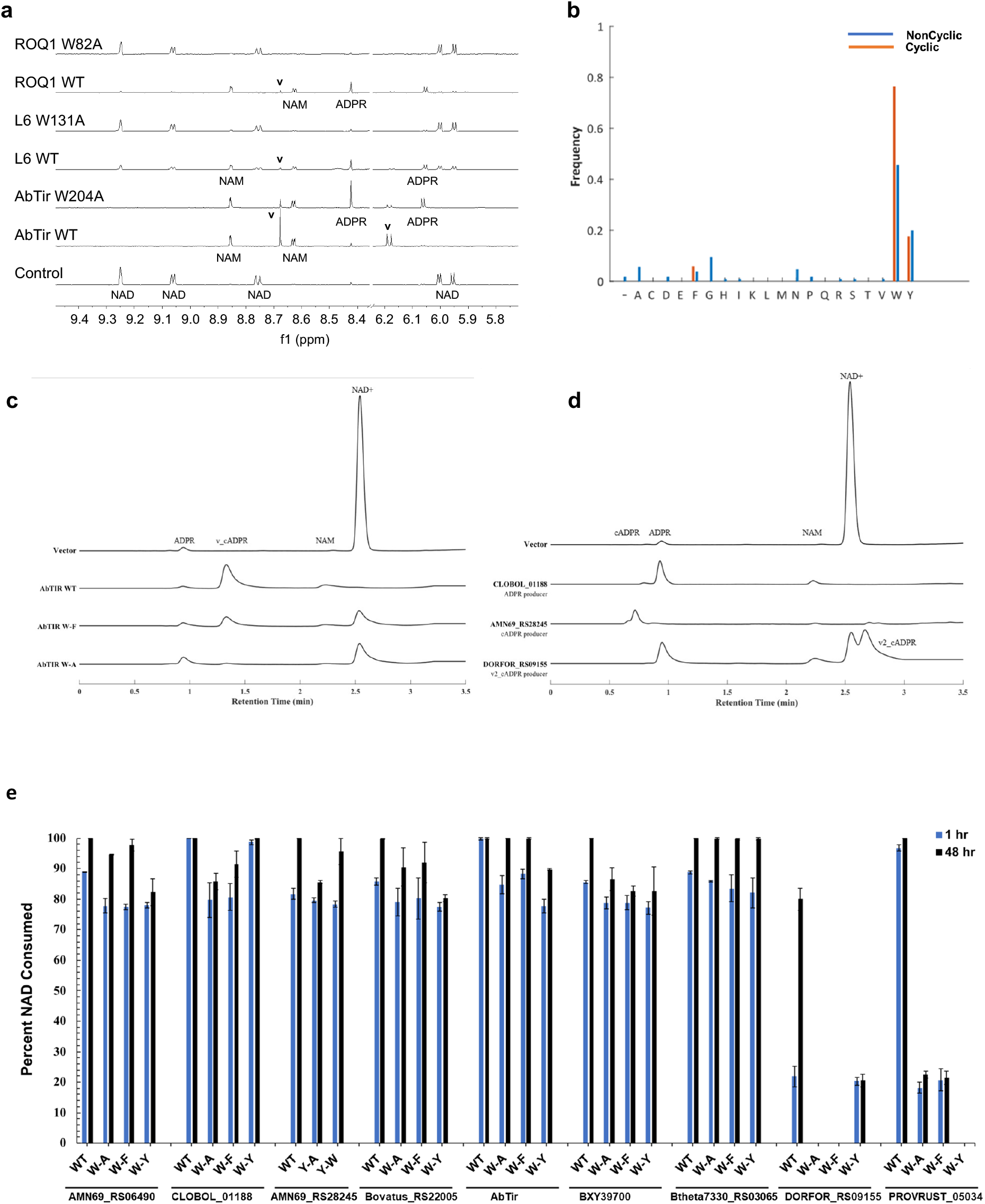
Conserved tryptophan is essential for ADPR cyclization. (a) Expansions of ^1^H NMR spectra showing altered NADase activity for AbTIR W204A, and absence of NADase activity for L6 W131A and ROQ1 W82A. The protein concentration was 50 μM for AbTir and 100 μM for L6 and ROQ1, while the initial NAD^+^ concentration was 500 μM. Spectra correspond to 24 h incubation time for AbTir samples and 16 h incubation time for L6 and ROQ1 samples. Selected peaks are labelled, showing the production of v-cADPR (**v**), Nam, and ADPR with WT proteins but not mutants. (b) Frequencies of amino-acids observed at the position equivalent to AbTir W204 in a multiple sequence alignment of 122 functionally characterized TIR domains. Each bar represents the frequency of the indicated amino-acid among TIR domains that do (cyclic, red) or do not (non-cyclic, blue) produce a cylic NAD^+^ catabolite, i.e. cADPR, v-cADPR (2’cADPR) or v2-cADPR (3’cADPR). (c-d) HPLC chromatograms of NAD^+^ consumption by different bacterial TIR domains. (c) HPLC chromatograms of metabolite extracts from wild-type and mutant AbTir reactions at 1 h. (d) HPLC chromatograms of metabolite extracts from various TIR domain reactions, illustrating the variety of NAD^+^ catabolites. (e) Percent of starting NAD^+^ consumed by wild-type and mutant TIR domains. Data are shown at 1 h and 48 h; n = 3 for all groups except where no data (ND) could be collected.

**Fig. S7.**
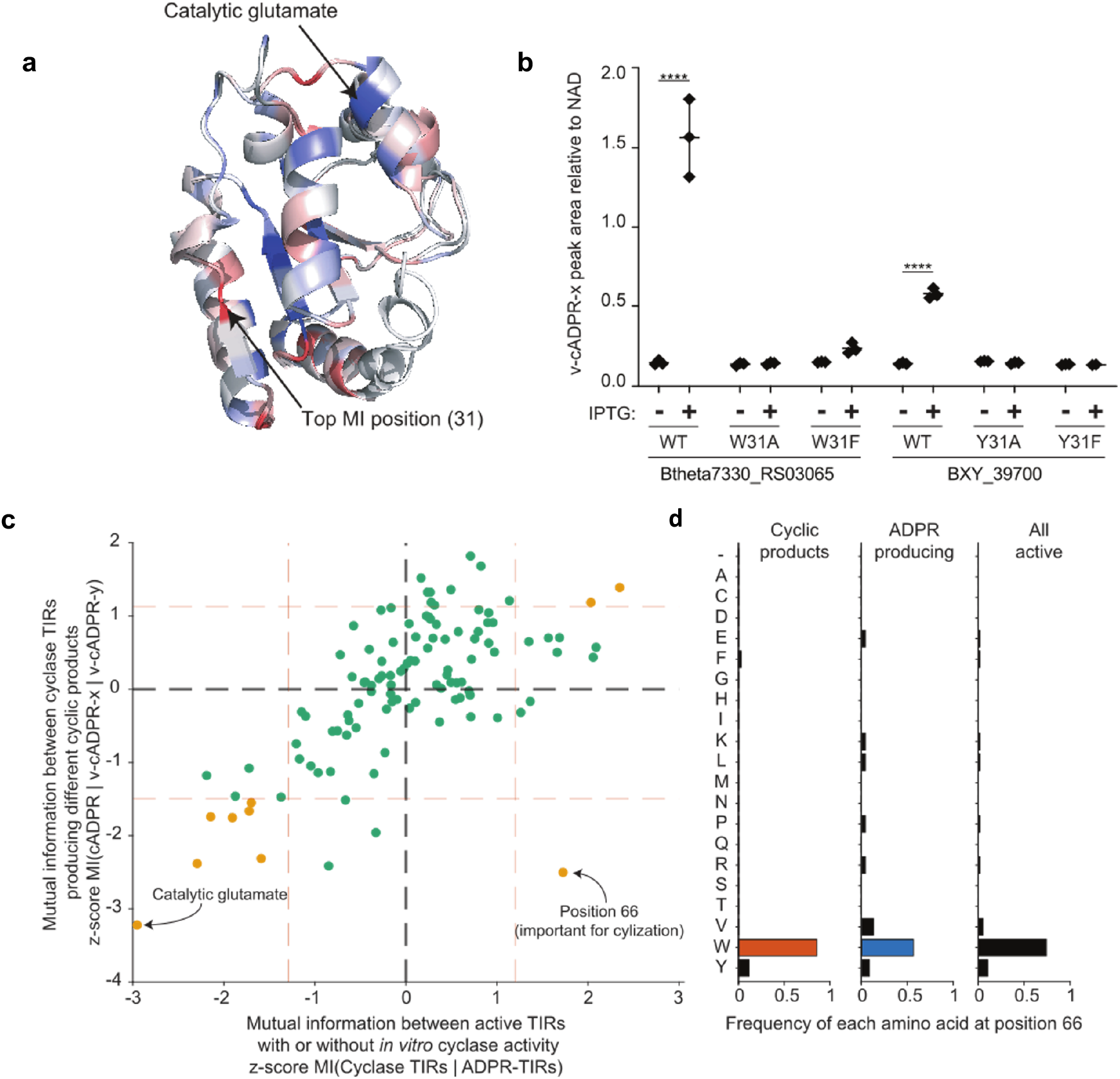
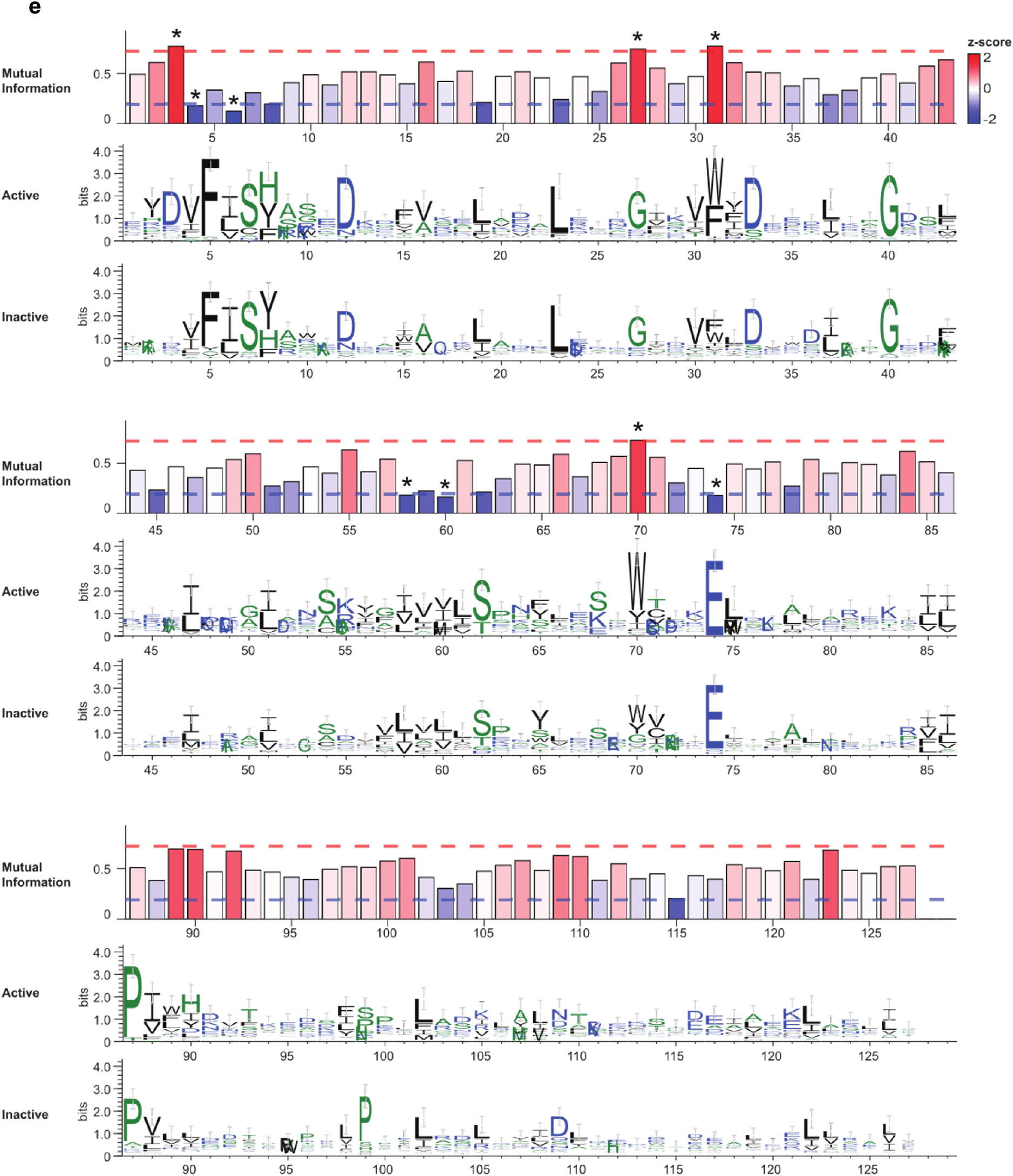
Mutual information analysis of TIR domains with, or without, NAD hydrolase activity in vitro. (a) Mutual information (MI) between ‘active’ (NADase-positive) and ‘inactive’ (NADase-negative) TIR domains (n = 40 and n = 70, respectively) illustrated on the superimposed ribbon structures of the v-cADPR (2’cADPR)-producing TIR domains from *Bacteroides xylanisolvens* XB1A and *Bacteroides thetaiotaomicron* 7330. These structures were modeled using BtpA from *Brucella melitensis* ATCC 23457 (PDB: 4LZP) as the template. Structures are colored based on the z-scored MI at each position, with red indicating positions that are most informative in delineating active from inactive TIR domains, and blue being the least informative. The top MI position (residue 31; tryptophan (W) in *B. thetaiotaomicron* 7330 and tyrosine (Y) in *B. xylanisolvens* XB1A), and the previously reported catalytic glutamate are shown. (b) Mutations at the position with the highest MI between active and inactive TIR domains in the v-cADPR (2’-cADPR) producing TIR domains from *B. thetaiotaomicron* 7330 (Btheta7330_RS03065) and *B. xylanisolvens* XB1A (BXY_39700). The peak area of v-cADPR (2’-cADPR) was normalized to the peak area of NAD^+^ measured in *E. coli* expressing wild-type (WT) and mutant TIR domains after a 1 h incubation in the presence or absence of the IPTG inducer of TIR expression (n=3). Note that endogenous NAD^+^ in *E. coli* served as the substrate for the TIR domains. (c) Scatter plot illustrating the relationship between positional mutual information calculated on an expanded set of TIR domain sequences (267 sequences, 110 positions following filtering) comparing (i) active TIR domains with or without in vitro cyclase activity [z-score MI(cyclase TIRs | ADPR-TIRs)] on the x-axis and (ii) between cyclase TIRs producing different cyclic products [z-score MI (cADPR | v-cADPR | v2-cADPR)] on the y-axis. Points corresponding to positions where the calculated MI was in the bottom or top 10th percentile (dashed red lines) for both sets of comparisons are colored in gold. The catalytic glutamate, which is fixed in both groups of sequences (MI = 0, lowest MI-Z-score) and position 66 in quadrant IV (corresponding to high MI between cyclase and ADPR producing TIRs and very low MI amongst TIRs that make different cyclic products) are both labeled. (d) Frequency of each amino acid at position 66 of the filtered alignment within sequences that produced cyclic products, ADPR, or all active sequences. (e) Mutual information (MI) identifies positions of importance to TIR enzymatic activity. MI calculated at 110 conserved positions in a multiple sequence alignment of TIR domains with and without cyclase activity. Bars are colored by the z-score of the MI. Red and blue dashed lines indicate MI thresholds (top and bottom 10th percentile, respectively) used to identify residues that are important for TIR enzymatic activity. Sequence logos generated from the multiple sequence alignment used to calculate MI between TIR domains that produced cyclic (n = 35) or non-cyclic (n = 23) products in vitro.

**Fig. S8.**
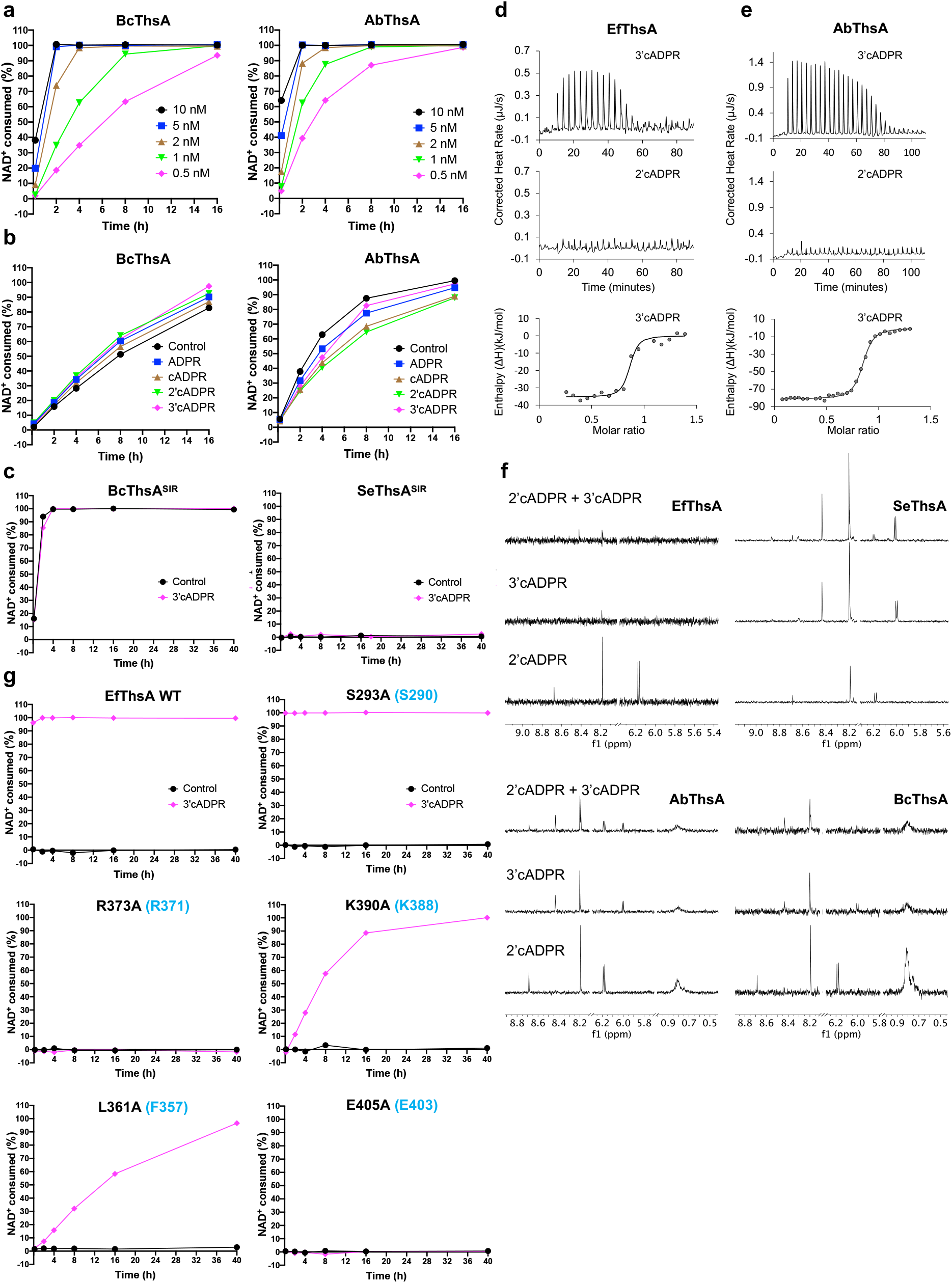
Characterization of ThsA NADase activity and cADPR isomer interaction by NMR and ITC. (a) NADase activity of of BcThsA (0.5 -10 nM μM) and AbThsA (0.5-10 nM). The initial NAD ^+^ concentration was 500 µM. (b) Activation of BcThsA (0.5 nM) and AbThsA (0.5 nM) NADase activity by 500 µM ADPR, cADPR, v-cADPR (2’cADPR) and v2-cADPR (3’cADPR). The initial NAD ^+^ concentration was 500 µM. (c) NADase activity of BcThsA^SIR2^ (0.5 nM) and SeThsA^SIR2^ (10 μM) in the absence and presence of 50 μM 3’cADPR. Initial NAD^+^ concentration was 500 μM. (d) Raw (top panel) and integrated (bottom panel) ITC data for the titration of 0.3 mM v2-cADPR (3’cADPR) with 35 µM EfThsA and raw ITC data for the titration of 0.3 mM v-cADPR (2’cADPR) with 35 µM EfThsA (middle panel). (e) Raw (top panel) and integrated (bottom panel) ITC data for the titration of 0.3 mM v2-cADPR (3’cADPR) with 50 µM AbThsA and raw ITC data for the titration of 0.3 mM v-cADPR (2’cADPR) with 50 µM AbThsA (middle panel). (f) STD NMR competition of v-cADPR (2’cADPR) vs v2-cADPR (3’cADPR) binding to EfThsA, SeThsA, AbThsA and BcThsA The protein concentration was 20 μM and the ligand concentration was 1 mM. (g) Effects of mutations on EfThsA (0.5 μM) NADase activity in the absence and presence of 50 μM v2-cADPR (3’cADPR). Initial NAD^+^ concentration was 500 μM. Corresponding residues in BcThsA are shown in blue.

**Fig. S9.**
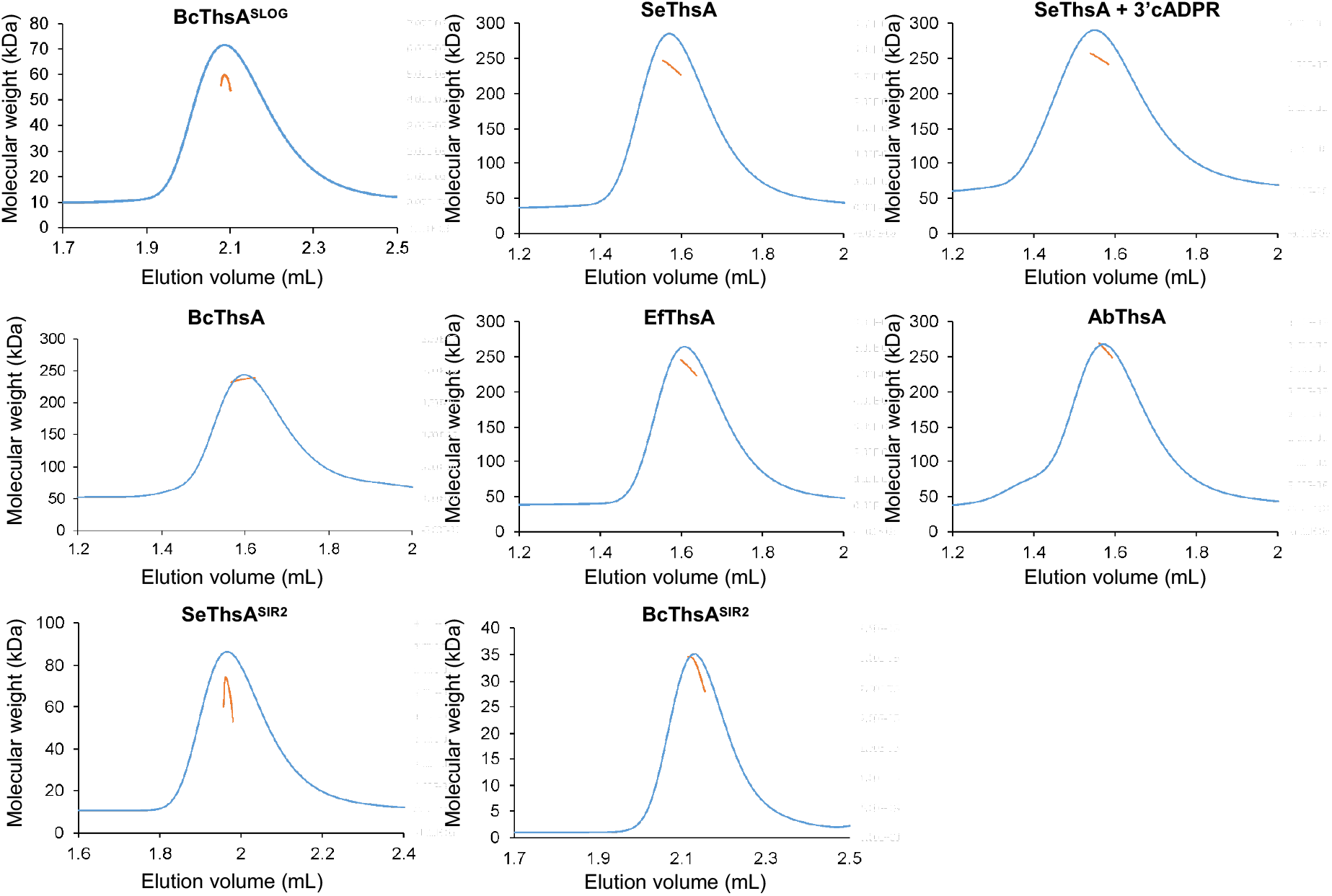
SEC-MALS analysis of ThsA proteins. The blue line represents the refractive index trace, while the orange line represents the average molecular mass distribution across the peak.

**Fig. S10.**
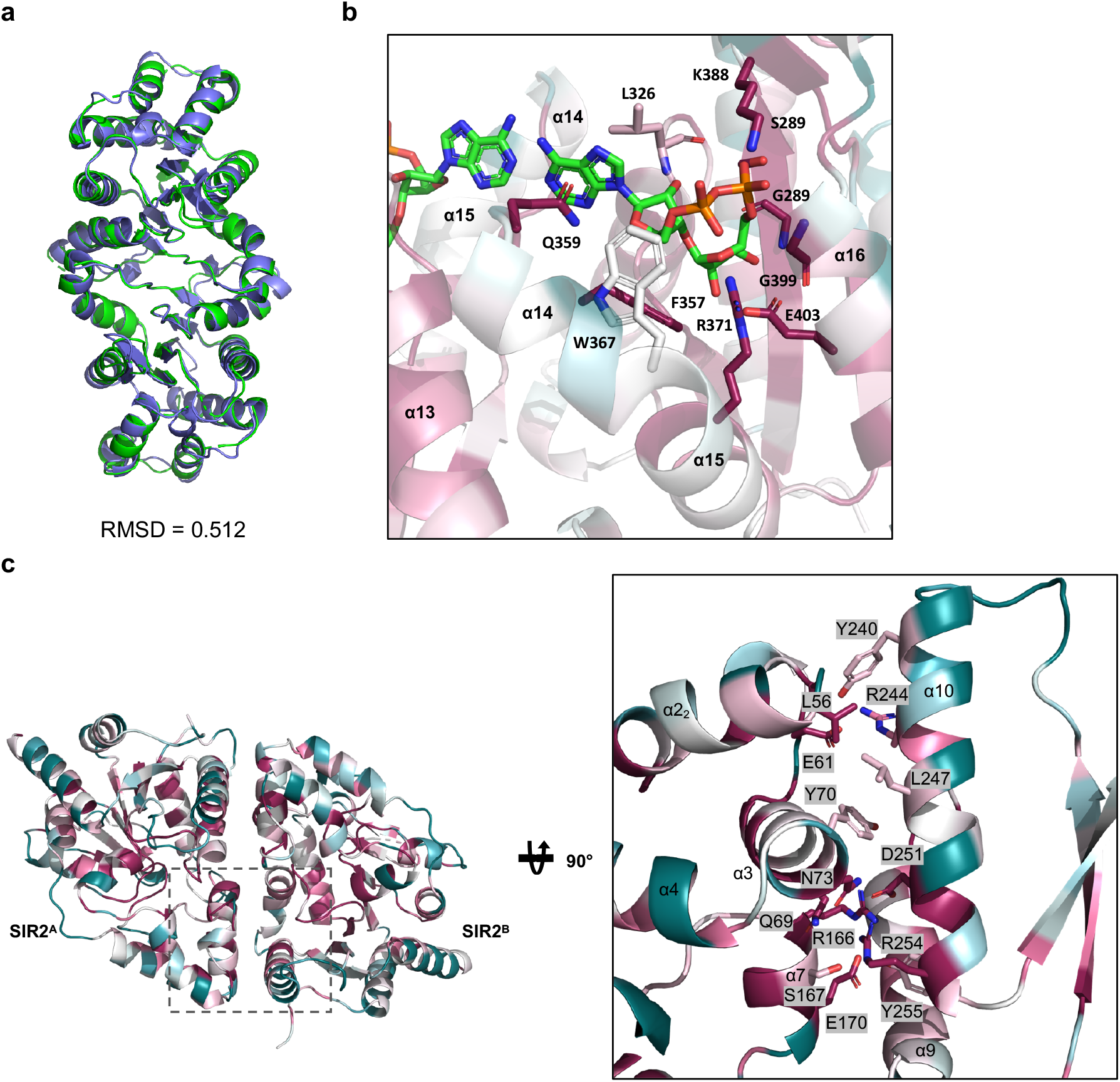
Structural analyses of BcThsA and SeThsA. (a) Structural superposition of BcThsA^SLOG^:v2-cADPR and BcThsA (PDB: 6LHX). (b) Enlarged cutaway of the v2-cADPR (3’cADPR)-binding pocket in the BcThsA^SLOG^ structure coloured by sequence conservation. Cyan corresponds to variable regions, while purple corresponds to conserved regions. Sequence conservation was calculated by ConSurf (*86*). (c) SeThsA SIR2 dimer coloured by sequence conservation. The insert shows an enlarged cutaway of ½ of the symmetric dimer interface with buried interface residues highlighted in stick representation.

**Fig. S11.**
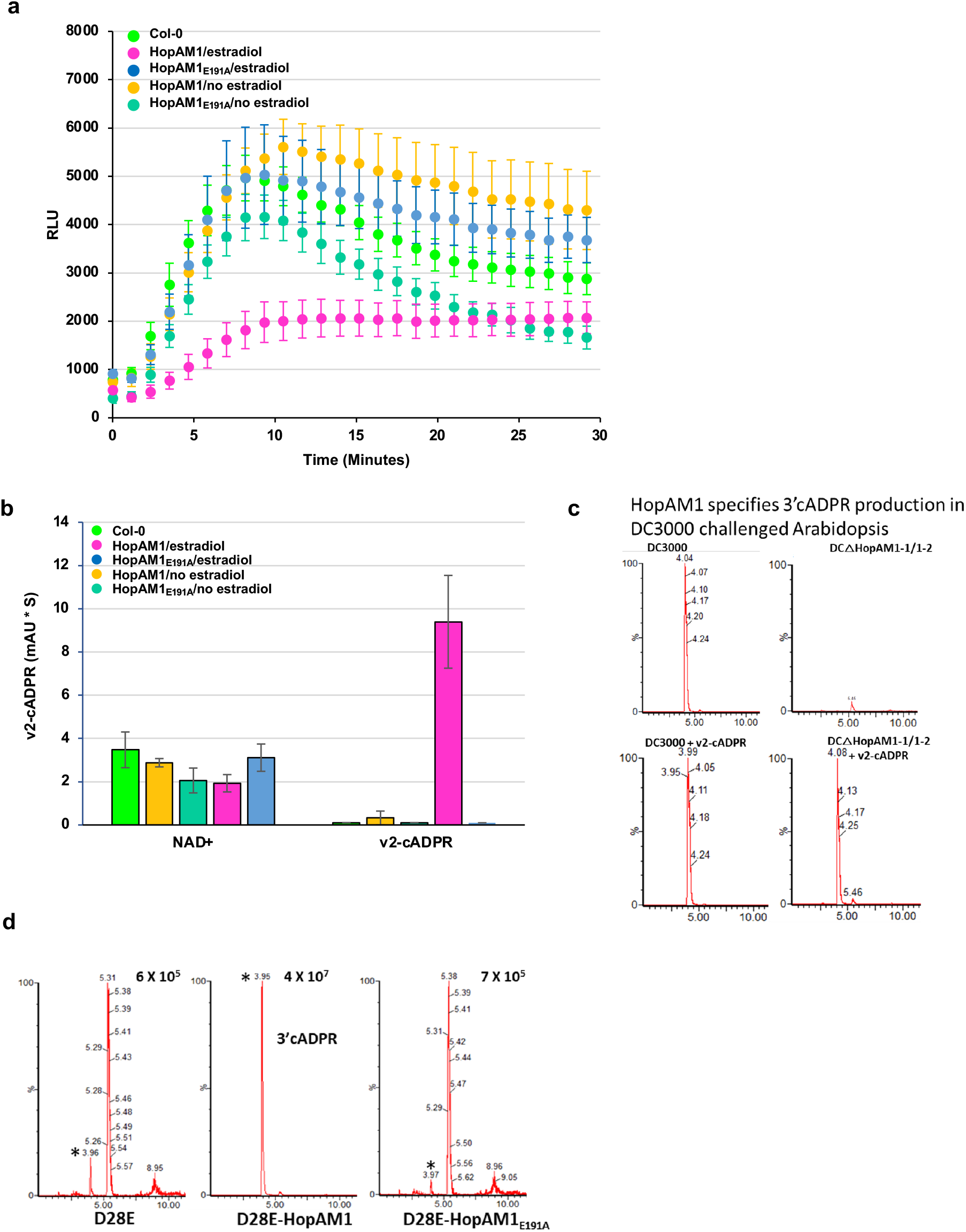
v2-cADPR production is associated with immunity suppression of *Pseudomonas syringae* DC3000 effector HopAM1. (a) ROS production of *Arabidopsis* transgenic plants expressing HopAM1 or HopAM1E191A induced with 10 mM estradiol. RLU, relative luminescence unit. ROS production is induced with the PAMP flg22 at 1 mM. (b) Quantification of NAD^+^ and v2-cADPR (3’cADPR) from transgenic leave samples in (b). (c) LC-MS/MS analysis of v2-cADPR (3’cADPR) production in *A. thaliana* Col-0 challenged with virulent *Pseudomonas syringae* pv. tomato strain DC3000 or DC3000 lacking both HopAM1-1 and HopAM1-2. Leaves were challenged with an inoculum of OD_600_ 0.15 and harvested 18 h post-infiltration, snap-frozen, freeze-dried and extracted in 10% methanol, 1% acetic acid. (d) LC-MS/MS analysis of 3’cADPR production in *A. thaliana* Col-0 challenged with the effector deficient DC3000 D28E strain carrying an empty vector, functional HopAM1 (D28E-HopAM1) or the catalytically inactive D28E-HopAM1^E191A^. Ion count for maximum peak shown, * corresponds to v2-cADPR (3’cADPR). Leaves were challenged with an inoculum of OD_600_ 0.15 and harvested 18 h post-infiltration, snap-frozen, freeze-dried and extracted in 10% methanol, 1% acetic acid.

**Table S1.**
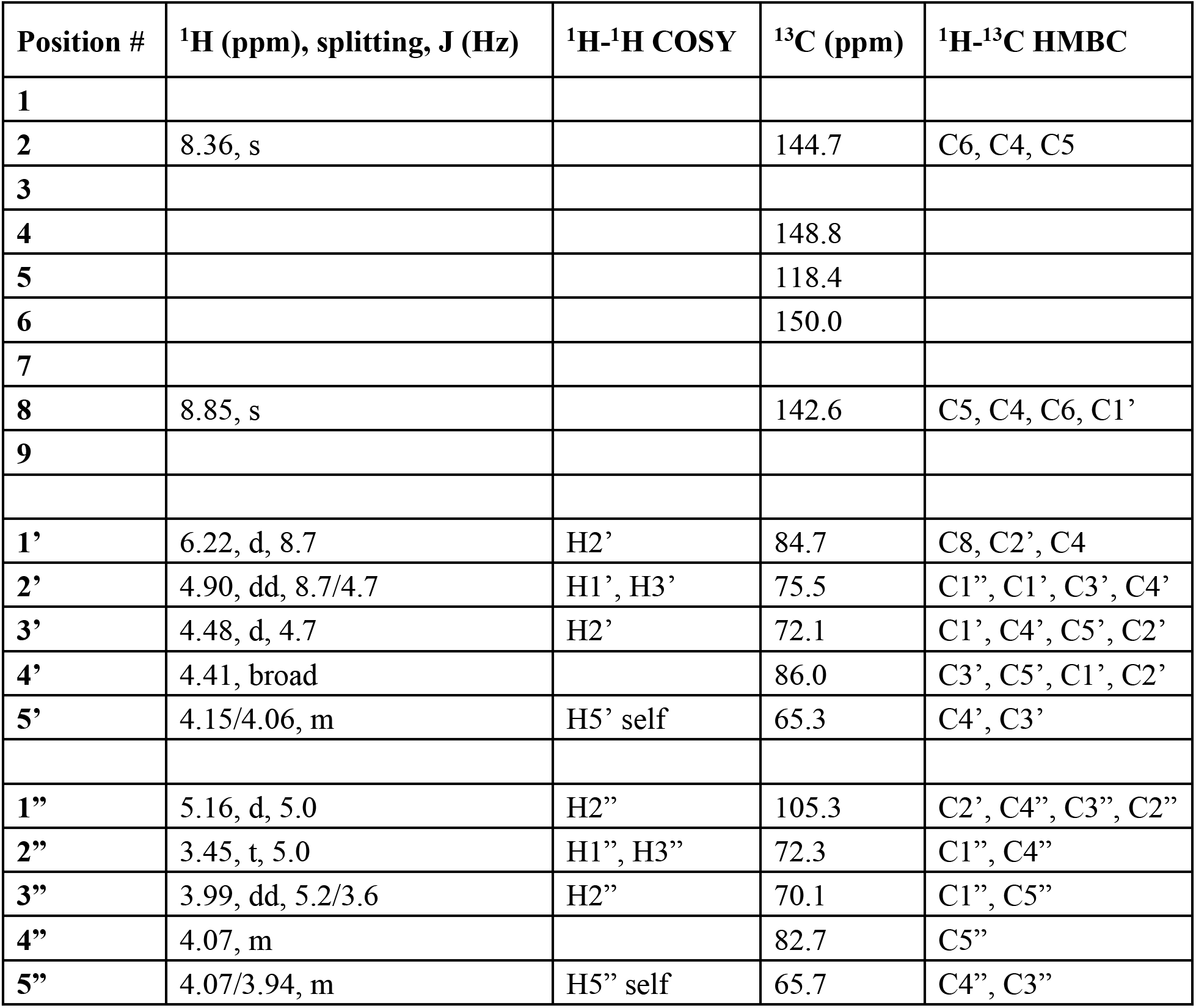
Assignments of v-cADPR (2’cADPR) NMR peaks (the structure shown in Figure 2).

| Position # | <sup>1</sup>H (ppm), splitting, J (Hz) | <sup>1</sup>H-<sup>1</sup>H COSY | <sup>13</sup>C (ppm) | <sup>1</sup>H-<sup>13</sup>C HMBC |
| --- | --- | --- | --- | --- |
| **1** |  |  |  |  |
| **2** | 8.36, s |  | 144.7 | C6, C4, C5 |
| **3** |  |  |  |  |
| **4** |  |  | 148.8 |  |
| **5** |  |  | 118.4 |  |
| **6** |  |  | 150.0 |  |
| **7** |  |  |  |  |
| **8** | 8.85, s |  | 142.6 | C5, C4, C6, C1' |
| **9** |  |  |  |  |
| **1'** | 6.22, d, 8.7 | H2' | 84.7 | C8, C2', C4 |
| **2'** | 4.90, dd, 8.7/4.7 | H1', H3' | 75.5 | C1'', C1', C3', C4' |
| **3'** | 4.48, d, 4.7 | H2' | 72.1 | C1', C4', C5', C2' |
| **4'** | 4.41, broad |  | 86.0 | C3', C5', C1', C2' |
| **5'** | 4.15/4.06, m | H5' self | 65.3 | C4', C3' |
| **1''** | 5.16, d, 5.0 | H2'' | 105.3 | C2', C4'', C3'', C2'' |
| **2''** | 3.45, t, 5.0 | H1'', H3'' | 72.3 | C1'', C4'' |
| **3''** | 3.99, dd, 5.2/3.6 | H2'' | 70.1 | C1'', C5'' |
| **4''** | 4.07, m |  | 82.7 | C5'' |
| **5''** | 4.07/3.94, m | H5'' self | 65.7 | C4'', C3'' |

**Table S2.**
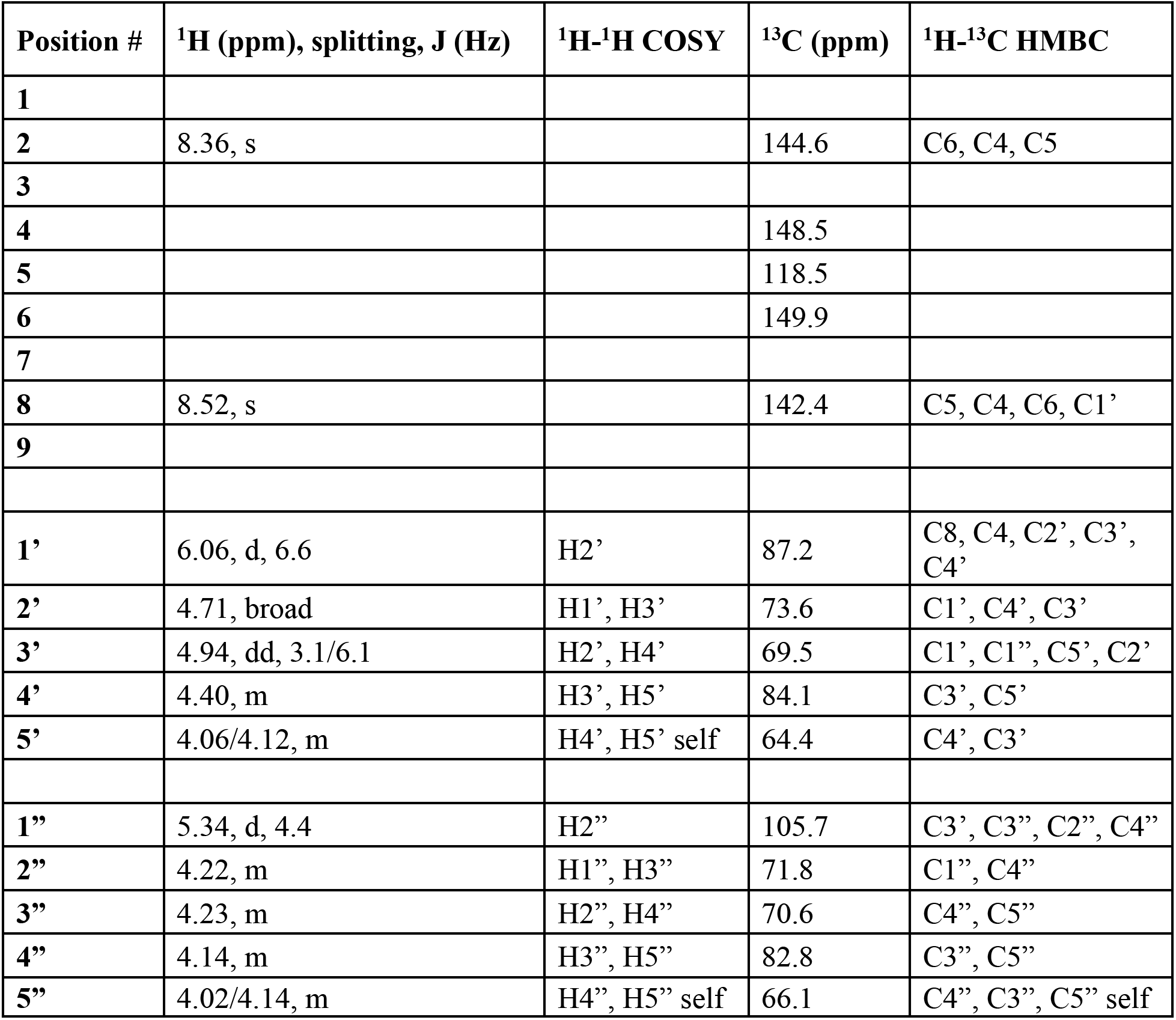
Assignments of v2-cADPR (3’cADPR) NMR peaks (the structure is shown in Figure 2).

**Table S3.** Assignments of NMR peaks of v2-cADPR (3’cADPR) purified from *N*. *benthamiana* leaves expressing the bacterial effector HopAM1.

| Position # | <sup>1</sup>H (ppm), splitting, J (Hz) | <sup>13</sup>C (ppm) | <sup>1</sup>H-<sup>13</sup>C HMBC | <sup>13</sup>C-<sup>31</sup>P J (Hz) |
| --- | --- | --- | --- | --- |
| 1 |  |  |  |  |
| 2 | 8.15, s | 153.0 | C4, C6 |  |
| 3 |  |  |  |  |
| 4 |  | 149.6 |  |  |
| 5 |  | 118.8 |  |  |
| 6 |  | 155.7 |  |  |
| 7 |  |  |  |  |
| 8 | 8.39, s | 139.6 | C4, C5 |  |
| 9 |  |  |  |  |
| 1' | 5.98, d, 7.7 | 85.6 | C2', C4, C8 |  |
| 2' | 4.74, broad | 72.8 | C1' |  |
| 3' | 4.93, dd, 1.2/5.9 | 69.4 | C1', C1'', C5' |  |
| 4' | 4.36, broad | 84.1 | C3', C5' | 9.4 |
| 5' | 4.09/3.97, m | 64.6 | C3', C4' | broad |
| 1'' | 5.35, d, 4.9 | 105.8 | C3', C4'' |  |
| 2'' | 4.23, m | 71.7 | C1'', C4'' |  |
| 3'' | 4.22, m | 70.7 | C1'', C5'' |  |
| 4'' | 4.14, m | 83.0 | C3'', C5'' | 9.6 |
| 5'' | 4.01/4.11, m | 66.1 | C3'', C4'' |  |

**Table S4.** Crystallographic data collection and refinement statistics.

|  | AbTir<sup>TIR</sup> | BtTir<sup>TIR</sup> | BcThsA<sup>SLOG:v2-</sup><br>cADPR | SeThsA |
| --- | --- | --- | --- | --- |
| **Data collection** |  |  |  |  |
| Space group | P 1 2<sub>1</sub> 1 | P 1 2<sub>1</sub> 1 | P 2<sub>1</sub> 2<sub>1</sub> 2<sub>1</sub> | C 2 2 2<sub>1</sub> |
| *a*, *b*, *c* (Å) | 54.68, 67.77,<br>72.17 | 37.69, 43.13,<br>98.76 | 46.19, 70.80,<br>121.74 | 98.81, 274.36,<br>89.54 |
| $\alpha$, $\beta$, $\gamma$ (°) | 90, 110.46, 90 | 90, 90.04, 90 | 90, 90, 90 | 90, 90, 90 |
| Resolution (Å) | 47.86 - 2.16<br>(2.24 - 2.16) | 43.13 - 1.42<br>(1.44 - 1.42) | 46.16 - 1.57 (1.59 -<br>1.57) | 92.96 - 3.40<br>(3.67 - 3.40) |
| Total reflections | 98447 (8184) | 407463<br>(17679) | 373868 (16957) | 104826<br>(22027) |
| Unique reflections | 25989 (2180) | 59368 (2685) | 56724 (2652) | 17169 (3473) |
| Completeness (%) | 97.90 (95.30) | 98.2 (88.8) | 99.7 (95.2) | 99.8 (99.9) |
| Multiplicity | 3.8 (3.8) | 6.9 (6.6) | 6.6 (6.4) | 6.1 (6.3) |
| Wilson B-factor<br>(Å<sup>2</sup>) | 25.33 | 15.59 | 13.84 | 97.29 |
| R-meas | 0.14 (0.77) | 0.07 (1.27) | 0.04 (0.26) | 0.12 (1.42) |
| R-merge | 0.12 (0.66) | 0.07 (1.17) | 0.04 (0.24) | 0.11 (1.31) |
| R-pim | 0.10 (0.52) | 0.03 (0.48) | 0.02 (0.10) | 0.05 (0.56) |
| Mean I/sigma(I) | 7.6 (1.7) | 11.2 (0.9) | 20.2 (4.6) | 6.7 (1.4) |
| CC<sub>1/2</sub> | 0.99 (0.77) | 0.99 (0.65) | 0.99 (0.97) | 0.99 (0.40) |
| **Refinement** |  |  |  |  |
| Resolution (Å) | 47.86 - 2.16 | 39.52-1.42 | 36.87 – 1.57 | 74.98 – 3.40 |
| Reflections used in<br>refinement | 25974 | 59326 | 56645 | 17149 |
| R-work | 0.1856 | 0.1890 | 0.1488 | 0.2351 |
| R-free | 0.2289 | 0.2085 | 0.1763 | 0.2912 |
| Number of non-<br>hydrogen atoms | 4653 | 2493 | 3606 | 7581 |
| Macromolecules | 4372 | 2328 | 3154 | 7559 |
| Ligands | 44 |  | 87 | 16 |
| Solvent | 237 | 165 | 365 |  |
| Protein residues | 545 | 285 | 390 | 936 |
| RMS bonds (Å) | 0.002 | 0.009 | 0.008 | 0.003 |
| RMS angles (°) | 0.62 | 1.05 | 0.945 | 0.641 |
| Ramachandran<br>favoured (%) | 96.83 | 96.70 | 96.63 | 94.85 |
| Ramachandran<br>allowed (%) | 3.17 | 3.30 | 3.37 | 5.04 |
| Ramachandran<br>outliers (%) | 0 | 0 | 0 | 0.11 |
| Rotamer outliers<br>(%) | 0.8 | 0 | 0 | 0 |
| Clash-score | 2.97 | 3.00 | 1.00 | 2.98 |
| Average B-factor<br>(Å<sup>2</sup>) | 29.66 | 27.19 | 17.84 | 145.61 |
| Macromolecules | 29.5 | 26.68 | 16.52 | 145.65 |
| Ligands | SO<sub>4</sub>: 39.86<br>PEG: 39.45 |  | 3'cADPR: 11.03<br>Glycerol: 24.56<br>SO<sub>4</sub>: 18.74 | PEG:123.34<br>Glycerol:<br>126.65 |
| Solvent | 30.38 | 34.33 | 29.22 |  |

The values in parentheses are for the highest-resolution shell. The statistics were calculated using Aimless (73) and MolProbability (78). $R_{\text{merge}} = \sum_{hkl} \sum_j |I_{hkl,j} - \langle I_{hkl} \rangle| / (\sum_{hkl} \sum_j I_{hkl,j})$ . $R_{\text{work}} / R_{\text{free}} = \sum_{hkl} |F_{hkl}^{\text{obs}} - F_{hkl}^{\text{calc}}| / (\sum_{hkl} F_{hkl}^{\text{obs}})$ ; $R_{\text{free}}$ was calculated using randomly chosen 3.5-10 % fraction of data that was excluded from refinement.

**Table S5.** CryoEM analysis of AbTir^TIR^:3AD

|  | AbTir<sup>TIR</sup>:3AD |
| --- | --- |
| **Data collection and processing** |  |
| Cryo-EM facility | University of Queensland |
| Microscope | JEOL CryoARM 300 |
| Detector | Gatan K3 |
| Voltage (kV) | 300 |
| Nominal magnification | 60,000 |
| Pixel size (Å) | 0.80 |
| Defocus range (µm) | -0.5 to -3.0 |
| Total exposure (e/Å<sup>2</sup>) | 40 |
| Exposure per frame (e/Å<sup>2</sup>) | 0.80 |
| Total micrographs (no.) | 2,019 |
| Total extracted particles (no.) | 730,607 |
| Final particles (no.) | 272,949 |
| Symmetry imposed | C1 |
| Map sharpening B-factor (Å<sup>2</sup>) | 127.6 |
| Resolution (FSC) |  |
| Masked (0.143) | 3.4 |
| Unmasked (0.143) | 4.1 |
| **Model composition** |  |
| Number of chains | 4 |
| Atoms | 8882 (hydrogens: 4374) |
| Residues | 536 |
| Water | 0 |
| Ligands | 3AD: 4 molecules |
| **Model validation** |  |
| Bonds (RMSD) |  |
| Length (Å) (> 4σ) | 0.002 (0) |
| Angles (°) (> 4σ) | 0.442 (0) |
| MolProbity score | 1.48 |
| Clash-score | 4.44 |
| Ramachandran plot (%) |  |
| Outliers | 0 |
| Allowed | 3.79 |
| Favored | 96.21 |
| Rotamer outliers (%) | 0 |
| Cβ≤ outliers (%) | 0 |
| Peptide plane (%) |  |
| Cis proline/general | 0.0/0.0 |
| Twisted proline/general | 0.0/0.0 |
| CaBLAM outliers (%) | 1.35 |
| ADP B-factor (min/max/mean; Å<sup>2</sup>) |  |
| Protein | 111.40/157.55/127.69 |
| Ligand | 119.10/127.44/122.90 |
| Occupancy = 1 (%) | 100 |
| Map to model FSC (0.143/0.5, Å) | 3.4/3.7 |
| Map correlation coefficient |  |
| Volume | 0.87 |
| Ligand (mean) | 0.80 |

The statistics were calculated using CryoSPARC (82) and the phenix.validation\_cryoem tool (87).

**Table S6.** Estimated changes of NAD^+^, ADPR, and 2’cADPR based on ^1^H NMR assays.

| AbTir | % change at 10 min |  |  | % change at 24 h |  |  |
| --- | --- | --- | --- | --- | --- | --- |
|  | NAD<sup>+</sup> | ADPR | 2’cADPR | NAD<sup>+</sup> | ADPR | 2’cADPR |
| WT | -100 | 0 | +100 | -100 | +1 | +99 |
| G174A | 0 | 0 | 0 | 0 | 0 | 0 |
| D175A | -100 | +3 | +97 | -100 | +3 | +97 |
| S176A | -2 | 0 | +2 | -8 | 0 | +8 |
| L177A | -89 | +13 | +76 | -99 | +15 | +84 |
| R178A<sup>1</sup> | -1 | 0 | +1 | -5 | +1 | +4 |
| D182A | -4 | 0 | +4 | -100 | +4 | +96 |
| W204A | -10 | +6 | +4 | -100 | +71 | +29 |
| T205A | -16 | +0 | +16 | -100 | +1 | +99 |
| Y207A | 0 | 0 | 0 | 0 | 0 | 0 |
| E208A<sup>2</sup> | 0 | 0 | 0 | -30<sup>3</sup> | 0<sup>3</sup> | +30<sup>3</sup> |
| E208D<sup>2</sup> | 0 | 0 | 0 | -8<sup>3</sup> | +8<sup>3</sup> | 0<sup>3</sup> |
| R215A | 0 | 0 | 0 | 0 | 0 | 0 |
| E216A | -100 | +5 | +95 | -100 | +5 | +95 |

<sup>1</sup> 30 μM protein
<sup>2</sup> 20 μM protein and 1 mM NAD<sup>+</sup>
<sup>3</sup> % change at 16 h

**Table S7.** MALS analysis of ThsA proteins.

| | MW (kDa)<br>Monomer<sup>1</sup> | MW (kDa)<br>Dimer<sup>1</sup> | MW (kDa)<br>Tetramer<sup>1</sup> | MW (kDa)<br>MALS | [Protein]<br>(μM)<sup>2</sup> |
| --- | --- | --- | --- | --- | --- |
| BcThsA | 57.9 | 115.7 | 231.5 | 236.1 +/- 0.5 | 30.0 |
| SeThsA | 57.0 | 114.1 | 228.2 | 237.9 +/- 18.3 | 17.5 |
| SeThsA + 3'cADPR | 57.0 | 114.1 | 228.2 | 250.6 +/- 2.5 | 17.5 |
| EfThsA | 57.7 | 115.5 | 231.0 | 235.5 +/- 3.5 | 17.3 |
| AbThsA | 58.1 | 116.2 | 232.3 | 260.3 +/- 2.6 | 17.2 |
| SeThsA<sup>SIR2</sup> | 33.4 | 66.7 | 133.4 | 60.0 +/- 6.2 | 30.0 |
| BcThsA<sup>SIR2</sup> | 33.2 | 66.5 | 132.9 | 32.3 +/- 1.1 | 30.1 |
| BcThsA<sup>SLOG</sup> | 24.8 | 49.6 | 99.2 | 51.6 +/- 4.1 | 80.7 |

<sup>1</sup>Calculated from amino acid sequence.
<sup>2</sup>Concentration of protein sample loaded onto SEC column.

**Table S8.** 3’cADPR-interacting residues in BcThsA, EfTshA, SeThsA and AbThsA, based on the BcThsA^SLOG^ crystal structure.

| BcThsA | EfThsA | SeThsA | AbThsA | Interacting moiety of 3'cADPR |
| --- | --- | --- | --- | --- |
| G289 | G292 | G288 | G287 | Distal ribose |
| S290 | S293 | S289 | S288 | Pyrophosphate |
| L326 | L330 | K325 | L324 | Adenine and adenine-linked ribose |
| F357 | L361 | F355 | F355 | Adenine-linked ribose |
| Q359 | L363 | Q357 | Q357 | Adenine and adenine-linked ribose |
| W367 | W369 | Y369 | W369 | Adenine-linked ribose |
| R371 | R373 | R373 | R373 | Pyrophosphate and distal ribose |
| K388 | K390 | K390 | K390 | Pyrophosphate |
| G399 | G401 | G398 | G407 | Pyrophosphate |
| E403 | E405 | E402 | E411 | Distal ribose |

## Notes

### Competing Interest Statement

The authors have declared no competing interest.

## References and Notes

1. T. Ve, S. J. Williams, B. Kobe, Structure and function of Toll/interleukin-1 receptor/resistance protein (TIR) domains. Apoptosis : an international journal on programmed cell death 20, 250–261 (2015).

2. A. Bowie, Translational mini-review series on Toll-like receptors: recent advances in understanding the role of Toll-like receptors in anti-viral immunity. Clin Exp Immunol 147, 217–226 (2007).

3. L. A. O’Neill, D. Golenbock, A. G. Bowie, The history of Toll-like receptors—redefining innate immunity. Nature Rev Immunol 13, 453–460 (2013).

4. S. Nimma, T. Ve, S. J. Williams, B. Kobe, Towards the structure of the TIR-domain signalosome. Curr Opin Struct Biol 43, 122–130 (2017).

5. K. Takeda, T. Kaisho, S. Akira, Toll-like receptors. Annual Rev Immunol 21, 335–376 (2003).

6. S. Nimma, W. Gu, N. Maruta, Y. Li, M. Pan, F. K. Saikot, B. Y. J. Lim, H. Y. McGuinness, Z. F. Zaoti, S. Li, S. Desa, M. K. Manik, J. D. Nanson, B. Kobe, Structural evolution of TIR-domain signalosomes. Front Immunol 12, 784484 (2021).

7. N. Maruta, H. Burdett, B. Y. J. Lim, X. Hu, S. Desa, M. K. Manik, B. Kobe, Structural basis of NLR activation and innate immune signalling in plants. Immunogenetics, (2022).

8. M. T. B. Clabbers, S. Holmes, T. W. Muusse, P. R. Vajjhala, S. J. Thygesen, A. K. Malde, D. J. B. Hunter, T. I. Croll, L. Flueckiger, J. D. Nanson, M. H. Rahaman, A. Aquila, M. S. Hunter, M. Liang, C. H. Yoon, J. Zhao, N. A. Zatsepin, B. Abbey, E. Sierecki, Y. Gambin, K. J. Stacey, C. Darmanin, B. Kobe, H. Xu, T. Ve, MyD88 TIR domain higher-order assembly interactions revealed by microcrystal electron diffraction and serial femtosecond crystallography. Nat Commun 12, 2578 (2021).

9. P. R. Vajjhala, T. Ve, A. Bentham, K. J. Stacey, B. Kobe, The molecular mechanisms of signaling by cooperative assembly formation in innate immunity pathways. Mol Immunol 86, 23–37 (2017).

10. T. Ve, P. R. Vajjhala, A. Hedger, T. Croll, F. DiMaio, S. Horsefield, X. Yu, P. Lavrencic, Z. Hassan, G. P. Morgan, A. Mansell, M. Mobli, A. O’Carroll, B. Chauvin, Y. Gambin, E. Sierecki, M. J. Landsberg, K. J. Stacey, E. H. Egelman, B. Kobe, Structural basis of TIR-domain-assembly formation in MAL- and MyD88-dependent TLR4 signaling. Nat Struct Mol Biol 24, 743–751 (2017).

11. M. Alaidarous, T. Ve, L. W. Casey, E. Valkov, D. J. Ericsson, M. O. Ullah, M. A. Schembri, A. Mansell, M. J. Sweet, B. Kobe, Mechanism of bacterial interference with TLR4 signaling by Brucella Toll/interleukin-1 receptor domain-containing protein TcpB. J Biol Chem 289, 654–668 (2014).

12. S. L. Chan, L. Y. Low, S. Hsu, S. Li, T. Liu, E. Santelli, G. Le Negrate, J. C. Reed, V. L. Woods, Jr., J. Pascual, Molecular mimicry in innate immunity: crystal structure of a bacterial TIR domain. J Biol Chem 284, 21386–21392 (2009).

13. C. Cirl, A. Wieser, M. Yadav, S. Duerr, S. Schubert, H. Fischer, D. Stappert, N. Wantia, N. Rodriguez, H. Wagner, C. Svanborg, T. Miethke, Subversion of Toll-like receptor signaling by a unique family of bacterial Toll/interleukin-1 receptor domain-containing proteins. Nat Med 14, 399–406 (2008).

14. J.-q. Fang, Q. Ou, J. Pan, J. Fang, D.-y. Zhang, M.-q. Qiu, Y.-q. Li, X.-H. Wang, X.-y. Yang, Z. Chi, TcpC inhibits toll-like receptor signaling pathway by serving as an E3 ubiquitin ligase that promotes degradation of myeloid differentiation factor 88. PLoS Pathog 17, e1009481 (2021).

15. P. R. Imbert, A. Louche, J. B. Luizet, T. Grandjean, S. Bigot, T. E. Wood, S. Gagne, A. Blanco, L. Wunderley, L. Terradot, P. Woodman, S. Garvis, A. Filloux, B. Guery, S. P. Salcedo, A Pseudomonas aeruginosa TIR effector mediates immune evasion by targeting UBAP1 and TLR adaptors. EMBO J 36, 1869–1887 (2017).

16. R. M. Newman, P. Salunkhe, A. Godzik, J. C. Reed, Identification and characterization of a novel bacterial virulence factor that shares homology with mammalian Toll/interleukin-1 receptor family proteins. Infect Immun 74, 594–601 (2006).

17. A. Waldhuber, M. Puthia, A. Wieser, C. Cirl, S. Dürr, S. Neumann-Pfeifer, S. Albrecht, F. Römmler, T. Müller, Y. Zheng, Uropathogenic Escherichia coli strain CFT073 disrupts NLRP3 inflammasome activation. J Clin Invest 126, 2425–2436 (2016).

18. D. Xiong, L. Song, S. Geng, Y. Jiao, X. Zhou, H. Song, X. Kang, Y. Zhou, X. Xu, J. Sun, Z. Pan, X. Jiao, Salmonella coiled-coil- and TIR-containing TcpS evades the Innate Immune system and subdues inflammation. Cell Rep 28, 804–818 e807 (2019).

19. M. Yadav, J. Zhang, H. Fischer, W. Huang, N. Lutay, C. Cirl, J. Lum, T. Miethke, C. Svanborg, Inhibition of TIR domain signaling by TcpC: MyD88-dependent and independent effects on Escherichia coli virulence. PLoS Pathog 6, e1001120 (2010).

20. J. D. Nanson, M. H. Rahaman, T. Ve, B. Kobe, Regulation of signaling by cooperative assembly formation in mammalian innate immunity signalosomes by molecular mimics. Semin Cell Dev Biol 99, 96–114 (2020).

21. S. Doron, S. Melamed, G. Ofir, A. Leavitt, A. Lopatina, M. Keren, G. Amitai, R. Sorek, Systematic discovery of antiphage defense systems in the microbial pangenome. Science 359, (2018).

22. B. R. Morehouse, A. A. Govande, A. Millman, A. F. A. Keszei, B. Lowey, G. Ofir, S. C. Shao, R. Sorek, P. J. Kranzusch, STING cyclic dinucleotide sensing originated in bacteria. Nature 586, 429-+ (2020).

23. N. Tal, B. R. Morehouse, A. Millman, A. Stokar-Avihail, C. Avraham, T. Fedorenko, E. Yirmiya, E. Herbst, A. Brandis, T. Mehlman, Y. Oppenheimer-Shaanan, A. F. A. Keszei, S. Shao, G. Amitai, P. J. Kranzusch, R. Sorek, Cyclic CMP and cyclic UMP mediate bacterial immunity against phages. Cell 184, 5728–5739 e5716 (2021).

24. B. Koopal, A. Potocnik, S. K. Mutte, C. Aparicio-Maldonado, S. Lindhoud, J. J. M. Vervoort, S. J. J. Brouns, D. C. Swarts, Short prokaryotic Argonaute systems trigger cell death upon detection of invading DNA. Cell 185, 1471–1486 e1419 (2022).

25. S. Horsefield, H. Burdett, X. Zhang, M. K. Manik, Y. Shi, J. Chen, T. Qi, J. Gilley, J. S. Lai, M. X. Rank, L. W. Casey, W. Gu, D. J. Ericsson, G. Foley, R. O. Hughes, T. Bosanac, M. von Itzstein, J. P. Rathjen, J. D. Nanson, M. Boden, I. B. Dry, S. J. Williams, B. J. Staskawicz, M. P. Coleman, T. Ve, P. N. Dodds, B. Kobe, NAD(+) cleavage activity by animal and plant TIR domains in cell death pathways. Science 365, 793–799 (2019).

26. L. Wan, K. Essuman, R. G. Anderson, Y. Sasaki, F. Monteiro, E. H. Chung, E. Osborne Nishimura, A. DiAntonio, J. Milbrandt, J. L. Dangl, M. T. Nishimura, TIR domains of plant immune receptors are NAD(+)-cleaving enzymes that promote cell death. Science 365, 799–803 (2019).

27. K. Essuman, D. W. Summers, Y. Sasaki, X. Mao, A. K. Y. Yim, A. DiAntonio, J. Milbrandt, TIR domain proteins are an ancient family of NAD(+)-consuming enzymes. Curr Biol 28, 421–430 e424 (2018).

28. K. Essuman, D. W. Summers, Y. Sasaki, X. Mao, A. DiAntonio, J. Milbrandt, The SARM1 Toll/interleukin-1 receptor domain possesses intrinsic NAD+ cleavage activity that promotes pathological axonal degeneration. Neuron 93, 1334–1343 (2017).

29. M. D. Figley, W. Gu, J. D. Nanson, Y. Shi, Y. Sasaki, K. Cunnea, A. K. Malde, X. Jia, Z. Luo, F. K. Saikot, T. Mosaiab, V. Masic, S. Holt, L. Hartley-Tassell, H. Y. McGuinness, M. K. Manik, T. Bosanac, M. J. Landsberg, P. S. Kerry, M. Mobli, R. O. Hughes, J. Milbrandt, B. Kobe, A. DiAntonio, T. Ve, SARM1 is a metabolic sensor activated by an increased NMN/NAD(+) ratio to trigger axon degeneration. Neuron 109, 1118–1136 e1111 (2021).

30. H. C. Lee, R. Aarhus, D. Levitt, The crystal structure of cyclic ADP-ribose. Nat Struct Biol 1, 143–144 (1994).

31. Z. Duxbury, S. Wang, C. I. MacKenzie, J. L. Tenthorey, X. Zhang, S. U. Huh, L. Hu, L. Hill, P. M. Ngou, P. Ding, Induced proximity of a TIR signaling domain on a plant-mammalian NLR chimera activates defense in plants. Proc Natl Acad Sci USA 117, 18832–18839 (2020).

32. J. M. Coronas-Serna, A. Louche, M. Rodríguez-Escudero, M. Roussin, P. R. Imbert, I. Rodríguez-Escudero, L. Terradot, M. Molina, J.-P. Gorvel, V. J. Cid, The TIR-domain containing effectors BtpA and BtpB from Brucella abortus impact NAD metabolism. PLoS Pathog 16, e1007979 (2020).

33. S. Eastman, T. Smith, M. A. Zaydman, P. Kim, S. Martinez, N. Damaraju, A. DiAntonio, J. Milbrandt, T. E. Clemente, J. R. Alfano, M. Guo, A phytobacterial TIR domain effector manipulates NAD(+) to promote virulence. New Phytol 233, 890–904 (2022).

34. G. Ofir, E. Herbst, M. Baroz, D. Cohen, A. Millman, S. Doron, N. Tal, D. B. A. Malheiro, S. Malitsky, G. Amitai, R. Sorek, Antiviral activity of bacterial TIR domains via immune signalling molecules. Nature 600, 116–120 (2021).

35. D. Ka, H. Oh, E. Park, J. H. Kim, E. Bae, Structural and functional evidence of bacterial antiphage protection by Thoeris defense system via NAD(+) degradation. Nat Commun 11, 2816 (2020).

36. A. M. Burroughs, D. Zhang, D. E. Schaffer, L. M. Iyer, L. Aravind, Comparative genomic analyses reveal a vast, novel network of nucleotide-centric systems in biological conflicts, immunity and signaling. Nucleic Acids Res 43, 10633–10654 (2015).

37. A. M. Burroughs, L. Aravind, Identification of uncharacterized components of prokaryotic immune systems and their diverse eukaryotic reformulations. J Bacteriol 202, (2020).

38. Y. Shi, P. S. Kerry, J. D. Nanson, T. Bosanac, Y. Sasaki, R. Krauss, F. K. Saikot, S. E. Adams, T. Mosaiab, V. Masic, X. Mao, F. Rose, E. Vasquez, M. Furrer, K. Cunnea, A. Brearley, W. Gu, Z. Luo, L. Brillault, M. J. Landsberg, A. DiAntonio, B. Kobe, J. Milbrandt, R. O. Hughes, T. Ve, Structural basis of SARM1 activation, substrate recognition, and inhibition by small molecules. Mol Cell 82, 1643–1659 e1610 (2022).

39. S. Ma, D. Lapin, L. Liu, Y. Sun, W. Song, X. Zhang, E. Logemann, D. Yu, J. Wang, J. Jirschitzka, Z. Han, P. Schulze-Lefert, J. E. Parker, J. Chai, Direct pathogen-induced assembly of an NLR immune receptor complex to form a holoenzyme. Science 370, (2020).

40. R. Martin, T. Qi, H. Zhang, F. Liu, M. King, C. Toth, E. Nogales, B. J. Staskawicz, Structure of the activated ROQ1 resistosome directly recognizing the pathogen effector XopQ. Science 370, (2020).

41. D. Yu, W. Song, E. Y. J. Tan, L. Liu, Y. Cao, J. Jirschitzka, E. Li, E. Logemann, C. Xu, S. Huang, A. Jia, X. Chang, Z. Han, B. Wu, P. Schulze-Lefert, J. Chai, TIR domains of plant immune receptors are 2′,3′-cAMP/cGMP synthetases mediating cell death. bioRxiv, 2021.2011.2009.467869 (2021).

42. B. Kaplan-Turkoz, T. Koelblen, C. Felix, M. P. Candusso, D. O’Callaghan, A. C. Vergunst, L. Terradot, Structure of the Toll/interleukin 1 receptor (TIR) domain of the immunosuppressive Brucella effector BtpA/Btp1/TcpB. FEBS Lett 587, 3412–3416 (2013).

43. G. A. Snyder, C. Cirl, J. Jiang, K. Chen, A. Waldhuber, P. Smith, F. Rommler, N. Snyder, T. Fresquez, S. Durr, N. Tjandra, T. Miethke, T. S. Xiao, Molecular mechanisms for the subversion of MyD88 signaling by TcpC from virulent uropathogenic Escherichia coli. Proc Natl Acad Sci U S A 110, 6985–6990 (2013).

44. J. S. Weagley, M. Zaydman, S. Venkatesh, Y. Sasaki, N. Damaraju, A. Yenkin, W. Buchser, D. A. Rodionov, A. Osterman, T. Ahmed, M. J. Barratt, A. DiAntonio, J. Milbrandt, J. I. Gordon, Products of gut microbial Toll/interleukin-1 receptor domain NADase activities in gnotobiotic mice and Bangladeshi children with malnutrition. Cell Rep 39, 110738 (2022).

45. Y. Xu, X. Tao, B. Shen, T. Horng, R. Medzhitov, J. L. Manley, L. Tong, Structural basis for signal transduction by the Toll/interleukin-1 receptor domains. Nature 408, 111–115 (2000).

46. A. Jia, S. Huang, W. Song, J. Wang, Y. Meng, Y. Sun, L. Xu, H. Laessle, J. Jirschitzka, J. Hou, T. Zhang, W. Yu, G. Hessler, E. Li, S. Ma, D. Yu, J. Gebauer, U. Baumann, X. Liu, Z. Han, J. Chang, J. E. Parker, J. Chai, TIR-catalyzed ADP-ribosylation reactions produce signaling molecules for plant immunity. bioRxiv, 2022.2005.2002.490369 (2022).

47. M. Howard, J. C. Grimaldi, J. F. Bazan, F. E. Lund, L. Santos-Argumedo, R. M. Parkhouse, T. F. Walseth, H. C. Lee, Formation and hydrolysis of cyclic ADP-ribose catalyzed by lymphocyte antigen CD38. Science 262, 1056–1059 (1993).

48. A. B. Bleecker, Genetic analysis of ethylene responses in *Arabidopsis thaliana*. Symp Soc Exp Biol 45, 149–158 (1991).

49. L. Aravind, D. Zhang, R. F. de Souza, S. Anand, L. M. Iyer, The natural history of ADP-ribosyltransferases and the ADP-ribosylation system. Curr Top Microbiol Immunol 384, 3–32 (2015).

50. K. Ueda, O. Hayaishi, ADP-ribosylation. Annu Rev Biochem 54, 73–100 (1985).

51. G. Hogrel, A. Guild, S. Graham, H. Rickman, S. Grushow, Q. Bertrand, L. Spagnolo, M. F. White, Cyclic nucleotide-induced superhelical structure activates a bacterial TIR immune effector. bioRxiv, 2022.2005.2004.490601 (2022).

52. G. A. Snyder, D. Deredge, A. Waldhuber, T. Fresquez, D. Z. Wilkins, P. T. Smith, S. Durr, C. Cirl, J. Jiang, W. Jennings, T. Luchetti, N. Snyder, E. J. Sundberg, P. Wintrode, T. Miethke, T. S. Xiao, Crystal structures of the Toll/Interleukin-1 receptor (TIR) domains from the Brucella protein TcpB and host adaptor TIRAP reveal mechanisms of molecular mimicry. J Biol Chem 289, 669–679 (2014).

53. A. Chaudhary, K. Ganguly, S. Cabantous, G. S. Waldo, S. N. Micheva-Viteva, K. Nag, W. S. Hlavacek, C. S. Tung, The Brucella TIR-like protein TcpB interacts with the death domain of MyD88. Biochem Biophys Res Commun 417, 299–304 (2012).

54. C. Felix, B. Kaplan Turkoz, S. Ranaldi, T. Koelblen, L. Terradot, D. O’Callaghan, A. C. Vergunst, The Brucella TIR domain containing proteins BtpA and BtpB have a structural WxxxE motif important for protection against microtubule depolymerisation. Cell Commun Signal 12, 53 (2014).

55. M. Bernoux, T. Ve, S. Williams, C. Warren, D. Hatters, E. Valkov, X. Zhang, J. G. Ellis, B. Kobe, P. N. Dodds, Structural and functional analysis of a plant resistance protein TIR domain reveals interfaces for self-association, signaling, and autoregulation. Cell Host Microbe 9, 200–211 (2011).

56. Q. Liu, R. Graeff, I. A. Kriksunov, H. Jiang, B. Zhang, N. Oppenheimer, H. Lin, B. V. Potter, H. C. Lee, Q. Hao, Structural basis for enzymatic evolution from a dedicated ADP-ribosyl cyclase to a multifunctional NAD hydrolase. J Biol Chem 284, 27637–27645 (2009).

57. R. Graeff, Q. Liu, I. A. Kriksunov, M. Kotaka, N. Oppenheimer, Q. Hao, H. C. Lee, Mechanism of cyclizing NAD to cyclic ADP-ribose by ADP-ribosyl cyclase and CD38. J Biol Chem 284, 27629–27636 (2009).

58. A. Leavitt, E. Yirmiya, G. Amitai, A. Lu, J. Garb, B. R. Morehouse, S. J. Hobbs, P. J. Kranzusch, R. Sorek, Viruses inhibit TIR gcADPR signaling to overcome bacterial defense. bioRxiv, 2022.2005.2003.490397 (2022).

59. T. Kurakawa, N. Ueda, M. Maekawa, K. Kobayashi, M. Kojima, Y. Nagato, H. Sakakibara, J. Kyozuka, Direct control of shoot meristem activity by a cytokinin-activating enzyme. Nature 445, 652–655 (2007).

60. T. Kuroha, H. Tokunaga, M. Kojima, N. Ueda, T. Ishida, S. Nagawa, H. Fukuda, K. Sugimoto, H. Sakakibara, Functional analyses of LONELY GUY cytokinin-activating enzymes reveal the importance of the direct activation pathway in Arabidopsis. Plant Cell 21, 3152–3169 (2009).

61. S. Huang, A. Jia, W. Song, G. Hessler, Y. Meng, Y. Sun, L. Xu, H. Laessle, J. Jirschitzka, S. Ma, Y. Xiao, D. Yu, J. Hou, R. Liu, H. Sun, X. Liu, Z. Han, J. Chang, J. E. Parker, J. Chai, Identification and receptor mechanism of TIR-catalyzed small molecules in plant immunity. bioRxiv, 2022.2004.2001.486681 (2022).

62. M. S. Drenichev, M. Bennett, R. A. Novikov, J. Mansfield, N. Smirnoff, M. Grant, S. N. Mikhailov, A role for 3′-O-β-D-ribofuranosyladenosine in altering plant immunity. Phytochemistry 157, 128–134 (2019).

63. M. Z. Li, S. J. Elledge, Harnessing homologous recombination in vitro to generate recombinant DNA via SLIC. Nat Meth 4, 251–256 (2007).

64. W. T. Mooij, E. Mitsiki, A. Perrakis, ProteinCCD: enabling the design of protein truncation constructs for expression and crystallization experiments. Nucleic Acids Res 37, W402–W405 (2009).

65. L. Stols, M. Gu, L. Dieckman, R. Raffen, F. R. Collart, M. I. Donnelly, A new vector for high-throughput, ligation-independent cloning encoding a tobacco etch virus protease cleavage site. Protein Expr Purif 25, 8–15 (2002).

66. F. W. Studier, Protein production by auto-induction in high-density shaking cultures. Protein Expr Purif 41, 207–234 (2005).

67. G. Pergolizzi, J. N. Butt, R. P. Bowater, G. K. Wagner, A novel fluorescent probe for NAD-consuming enzymes. Chem Commun 47, 12655–12657 (2011).

68. M. Piotto, V. Saudek, V. Sklenar, Gradient-tailored excitation for single-quantum NMR spectroscopy of aqueous solutions. J Biomol NMR 2, 661–665 (1992).

69. V. Sklenar, M. Piotto, R. Leppik, V. Saudek, Gradient-tailored water suppression for 1H- 15N HSQC experiments optimized to retain full sensitivity. J Magn Reson 102, 241–245 (1993).

70. M. Mayer, B. Meyer, Characterization of ligand binding by saturation transfer difference NMR spectroscopy. Angew Chem Int Ed Engl 38, 1784–1788 (1999).

71. A. Chaikuad, S. Knapp, F. von Delft, Defined PEG smears as an alternative approach to enhance the search for crystallization conditions and crystal-quality improvement in reduced screens. Acta Crystallogr D Biol Crystallogr 71, 1627–1639 (2015).

72. W. Kabsch, Xds. Acta Crystallogr D BiolCrystallogr 66, 125–132 (2010).

73. P. Evans, Scaling and assessment of data quality. Acta Crystallogr D Biol Crystallogr 62, 72–82 (2006).

74. P. D. Adams, D. Baker, A. T. Brunger, R. Das, F. DiMaio, R. J. Read, D. C. Richardson, J. S. Richardson, T. C. Terwilliger, Advances, interactions, and future developments in the CNS, Phenix, and Rosetta structural biology software systems. Ann Rev Biophys 42, 265–287 (2013).

75. P. Emsley, B. Lohkamp, W. G. Scott, K. Cowtan, Features and development of Coot. Acta Crystallogr D Biol Crystallogr 66, 486–501 (2010).

76. A. J. McCoy, Solving structures of protein complexes by molecular replacement with Phaser. Acta Crystallogr D 63, 32–41 (2007).

77. P. V. Afonine, R. W. Grosse-Kunstleve, N. Echols, J. J. Headd, N. W. Moriarty, M. Mustyakimov, T. C. Terwilliger, A. Urzhumtsev, P. H. Zwart, P. D. Adams, Towards automated crystallographic structure refinement with phenix.refine. Acta Crystallogr D 68, 352–367 (2012).

78. V. B. Chen, W. B. Arendall, 3rd, J. J. Headd, D. A. Keedy, R. M. Immormino, G. J. Kapral, L. W. Murray, J. S. Richardson, D. C. Richardson, MolProbity: all-atom structure validation for macromolecular crystallography. Acta Crystallogr D Biol Crystallogr 66, 12–21 (2010).

79. T. G. Battye, L. Kontogiannis, O. Johnson, H. R. Powell, A. G. Leslie, iMOSFLM: a new graphical interface for diffraction-image processing with MOSFLM. Acta Crystallogr D Biol Crystallogr 67, 271–281 (2011).

80. J. Jumper, R. Evans, A. Pritzel, T. Green, M. Figurnov, O. Ronneberger, K. Tunyasuvunakool, R. Bates, A. Zidek, A. Potapenko, A. Bridgland, C. Meyer, S. A. A. Kohl, A. J. Ballard, A. Cowie, B. Romera-Paredes, S. Nikolov, R. Jain, J. Adler, T. Back, S. Petersen, D. Reiman, E. Clancy, M. Zielinski, M. Steinegger, M. Pacholska, T. Berghammer, S. Bodenstein, D. Silver, O. Vinyals, A. W. Senior, K. Kavukcuoglu, P. Kohli, D. Hassabis, Highly accurate protein structure prediction with AlphaFold. Nature 596, 583–589 (2021).

81. T. I. Croll, ISOLDE: a physically realistic environment for model building into low- resolution electron-density maps. Acta Crystallogr D Struct Biol 74, 519–530 (2018).

82. A. Punjani, J. L. Rubinstein, D. J. Fleet, M. A. Brubaker, cryoSPARC: algorithms for rapid unsupervised cryo-EM structure determination. Nat Methods 14, 290–296 (2017).

83. E. F. Pettersen, T. D. Goddard, C. C. Huang, E. C. Meng, G. S. Couch, T. I. Croll, J. H. Morris, T. E. Ferrin, UCSF ChimeraX: Structure visualization for researchers, educators, and developers. Protein Sci 30, 70–82 (2021).

84. P. V. Afonine, B. K. Poon, R. J. Read, O. V. Sobolev, T. C. Terwilliger, A. Urzhumtsev, P. D. Adams, Real-space refinement in PHENIX for cryo-EM and crystallography. Acta Crystallogr D Struct Biol 74, 531–544 (2018).

85. S. Asai, K. Ohta, H. Yoshioka, MAPK signaling regulates nitric oxide and NADPH oxidase-dependent oxidative bursts in *Nicotiana benthamiana*. Plant Cell 20, 1390–1406 (2008).

86. H. Ashkenazy, S. Abadi, E. Martz, O. Chay, I. Mayrose, T. Pupko, N. Ben-Tal, ConSurf 2016: an improved methodology to estimate and visualize evolutionary conservation in macromolecules. Nucleic Acids Res 44, W344–350 (2016).

87. C. J. Williams, J. J. Headd, N. W. Moriarty, M. G. Prisant, L. L. Videau, L. N. Deis, V. Verma, D. A. Keedy, B. J. Hintze, V. B. Chen, S. Jain, S. M. Lewis, W. B. Arendall, 3rd, J. Snoeyink, P. D. Adams, S. C. Lovell, J. S. Richardson, D. C. Richardson, MolProbity: More and better reference data for improved all-atom structure validation. Protein Sci 27, 293–315 (2018).

